# Dehydrozingerone mitigates energy deficits and cognitive impairments induced by cranial irradiation

**DOI:** 10.64898/2026.05.06.723293

**Authors:** Anuradha Kesharwani, PavanKalyan Banavath, Akuthota Akanksha, Richa Chauhan, Vinita Trivedi, Krishna Pandey, Velayutham Ravichandiran, Vipan Kumar Parihar

## Abstract

Radiotherapy is widely used in the management of brain tumors; however, it is often associated with delayed adverse effects, including cognitive decline and depression-like behavior. These effects are thought to arise, in part, from suppressed hippocampal neurogenesis, altered neuronal architecture, and microglial dysfunction. Despite this, the precise mechanisms underlying irradiation-induced cognitive deficits, as well as effective therapeutic interventions, remain poorly understood. In the present study, six-month-old male mice were subjected to a single 9 Gy dose of cranial irradiation, followed by behavioral assessments several weeks post-exposure. We observed that cranial irradiation significantly impaired hippocampal-, prefrontal cortex-, and cortical-dependent memory functions. Notably, treatment with dehydrozingerone (DH), a curcumin analog (50 mg/kg, oral administration for two weeks), markedly prevented these cognitive deficits. At the molecular level, irradiation disrupted the activity of key enzymes involved in the tricarboxylic acid (TCA) cycle and the glutamate–glutamine/GABA cycle, both of which were restored following DH treatment. Furthermore, irradiation induced dysregulation of genes and proteins associated with glycolysis (Atp2b1, mt-Nd2, mt-Atp6), mitochondrial energetics (mt-Atp8, mt-Cytb), glucose transport (Slc4a5), insulin resistance (Etnppl), lipid metabolism (Pla2g3, Plin4), and inflammation (Ighg2c), all of which were significantly normalized by DH. Importantly, DH also prevented irradiation-induced loss of cell-type-specific glucose transporter expression, including GLUT3 in neurons and GLUT5 in microglia. In conclusion, our findings suggest that DH is a promising therapeutic candidate for mitigating irradiation-induced energy deficits and cognitive impairments, likely through modulation of metabolic and mitochondrial pathways.

## Introduction

Radiotherapy is a standard treatment for CNS malignancies and remains a first-line therapy for most paediatric and adult brain cancers. While cranial irradiation significantly slows the progression of brain cancer and increases survival rates, it has also resulted in an increase in the proportion of long-term survivors with severe neurocognitive impairment. Several studies have reported that cranial irradiation associated cognitive deficits, involves altered hippocampal neurogenesis, chronic inflammation, increased oxidative stress, white matter disruption, and long-term changes in cerebral blood flow and metabolism (Makale et al., 2017; Owonikoko et al., 2014). These patterns of damage are characteristic features of CNS radiation injury and may alter energy metabolism and mitochondrial function in the brain, contributing to progressive cognitive dysfunction.

Aberrant energy metabolism has been widely implicated in the pathogenesis of type II diabetes mellitus, cancer, aging, and organ fibrosis. However, its link on radiation-induced cognitive deficits has not been fully explored. Energy metabolism is essential for the production of ATP, amino acids, and fatty acid intermediates that drive biosynthetic processes involved in cells and tissue restoration. The brain is an energy-intensive organ that depends on glucose as its main energy source, and tight regulation of glucose metabolism is critical for normal function; disturbances in this process can contribute to neurological disorders such as Alzheimer disease (AD) progression and cognitive impairments (Mergenthaler et al., 2013). Therefore, elucidating the impact of radiation on energy metabolism and mitochondrial energetics is essential for understanding the underlying basis, which may uncover a new mechanism for restoring or preventing the cognitive deficits in patients receiving radiotherapy.

Here, we show that cranial irradiation alters the cognitive function, in part by affecting the energy metabolism, mitochondrial energetics, glucose transporters and disrupting neurotransmitter imbalance. Furthermore, we demonstrate that these radiation-associated deficits can be reversed by dehydrozingerone (DH), a curcumin analogue with antioxidant and neuroprotective properties. This study links the energy metabolism and mitochondrial energetics as major deficits by radiation that can reverses by DH intervention.

## Methods

### Animals and Irradiation

All animal studies in this study were performed in compliance with the guidelines established by the Committee for Control and Supervision of Experiments on Animals (CPCSEA), Ministry of Environment and Forests, Government of India, New Delhi. The experimental protocols were approved by the Institutional Animal Ethics Committee of the National Institute of Pharmaceutical Education and Research, Hajipur, India (844102). Male C57BL/6J mice, six months of age, were obtained from the ICMR National Institute of Nutrition, Hyderabad, India. The animals were housed at the Central Animal Research Facility of NIPER Hajipur under controlled environmental conditions, including a temperature of 20 ± 1 °C, relative humidity of 50 ± 10%, and a 12:12 h light–dark cycle. Six-month-old wild-type male C57BL/6J mice were selected for this study, as this age approximates the stage in humans when the incidence of glioblastoma multiforme (GBM) is most common, with a median diagnosis age of approximately 64 years. The mice were randomly assigned to four experimental groups (n = 10 per group): unirradiated mice receiving saline as vehicle (0 Gy + Veh), unirradiated mice receiving DH (0 Gy + DH), irradiated mice receiving saline (9 Gy + Veh), and irradiated mice receiving DH (9 Gy + DH). Cranial irradiation was performed under anaesthesia using 5% isoflurane for induction and 2% for maintenance (vol/vol). Animals were positioned ventrally on the treatment platform of a cobalt irradiator to deliver head-only irradiation at a dose rate of 1.0 Gy/min (Parihar and Limoli, 2013). One week following cranial irradiation, mice in the DH treatment group were administered DH orally at a dose of 50 mg/kg for two weeks. After completion of the treatment period, animals underwent a series of behavioral tests to evaluate mood- and memory-related functions. These included the Novel Object Recognition (NOR), Object-in-Place (OiP), Temporal Order (TO), water maze, and Forced Swim tests (***Figure 1***). To minimize circadian variability, all behavioral experiments were conducted during the light phase between 9:00 and 14:00 hours. Detailed protocols for the behavioral assessments are provided in ***Supplementary File 1***.

**Fig. 1:**
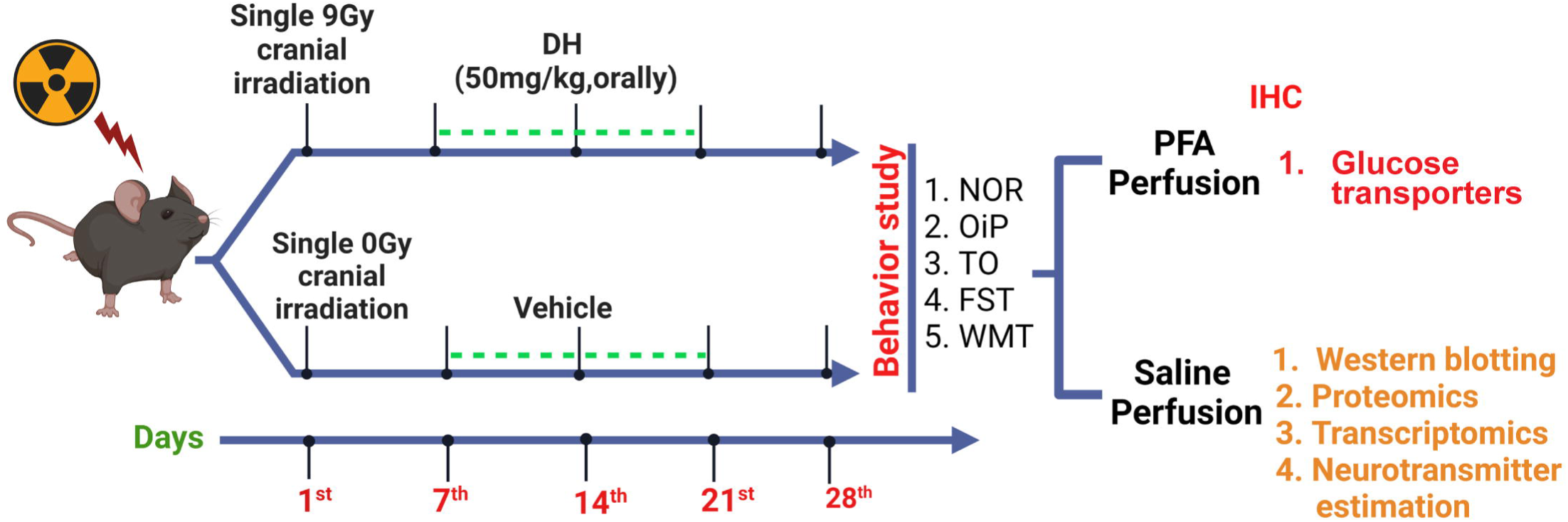
The diagram presents the series of in vivo experiments and analyses conducted in the study. The experimental protocol consisted of cranial irradiation utilizing a cobalt irradiator, succeeded by the administration of DH treatment. Animals were euthanized 24 hours following the final behavioral assessment. Brains were then collected and preserved at -80°C for future biochemical analyses.

### Behavioral Analysis

The Novel Object Recognition (NOR) test evaluates the preference for novelty, which primarily depends on the proper functioning of the medial prefrontal cortex (mPFC). In contrast, the Object-in-Place (OiP) task assesses associative recognition memory, which relies on interactions between distinct brain regions, particularly the hippocampus, perirhinal cortex, and mPFC (Barker and Warburton, 2015; Kesharwani et al., 2025a; Kesharwani et al., 2025b; Kesharwani, 2026). The Temporal Order (TO) measures the capacity to distinguish previously explored objects based on their relative recency, reflecting the capacity to recall past events. This task specifically relies on the crosstalk between the perirhinal cortex and mPFC (Barker and Warburton, 2015). All objects used in this study were selected to ensure equal inherent preference and were easily distinguishable by control mice. To maintain consistency, the objects were kept at a constant height and volume, but varied in shape, texture, and appearance. During behavioral testing, objects were randomly assigned to reduce any potential object-related bias on preference behavior. The Morris water maze test is used to evaluate spatial learning and memory in mice. This task relies primarily on the functional integrity of the hippocampus, a key brain region involved in spatial navigation and memory formation (Kodali et al., 2015). Data were analysed by two experimenters who were unaware of the treatment groups of the animals. A discrimination index (DI) was calculated to quantify novelty preference. A positive DI score indicates a preference for novelty, while a negative DI score suggests indifference or reduced interest in novelty (Kesharwani et al., 2025a; Kesharwani et al., 2025b; Kesharwani, 2026; Parihar et al., 2020; Parihar et al., 2021).

### Processing of biological tissues and immunohistochemical analysis

Twenty-four hours after the final behavioral test, the mice were euthanized under deep anaesthesia, and their brains were harvested for biochemical and immunohistochemical analyses (Christie et al., 2012; Kesharwani et al., 2025a; Kesharwani et al., 2025b; Kesharwani, 2026). As part of the biochemical estimation process, the mice underwent a procedure and then perfused using chilled heparinized saline Following this, their brains were promptly isolated from the skull. The hippocampus was meticulously isolated from the brain and utilized for transcriptomics, proteomics, and neurotransmitter analysis. For immunohistochemical analysis, mice were perfused with saline and then 4% paraformaldehyde (4% PFA) was used. ***Supplementary File 1*** provides a comprehensive overview of the methodologies used for transcriptome, proteome, and immunohistochemistry analysis.

### RNA Sequencing

RNA sequencing was conducted on saline perfused hippocampus of mice brains at the LCGC Wipro Genome Centre in Bengaluru. Tissue RNA was quantified using the RNeasy small kit (Cat no. 74106). The data from RNA-seq were standardised with the UCSC Genome Browser on mouse (version: mm39, GRCm39 (GCA_000001635.9), Jun. 2020) with the STAR splice aware alignment tool version 2.7.6a prior to the transcriptomics sequencing investigation, which was carried out with Illumina Nextseq 2000 and Fastqc. The quality of RNA was assessed using the Tapestation 4200™ (Agilent Technologies, Santa Clara, US), and the NanoDrop ND-100 Spectrophotometer (NanoDrop Technologies, Wilmington, DE) and Qubit 4 Fluorometer (Thermo Fisher Scientific, IN) were used to quantify RNA concentration (Dobin et al., 2013). The three most exceptional RNA samples from each cohort were selected, assigned unique identifiers, and sequenced. The Illumina Nextseq 2000 and Fastqc was used for transcriptomics sequencing study. Genes showing differential expression were identified with a false discovery rate (FDR) of 0.05, corresponding to a 95% probability of true differential expression, and an absolute log□fold-change of by comparing individual groups to unirradiated mice (0 Gy + Veh). The microarray was employed to elucidate the altered gene expression, and the differential expression of each gene was identified (Kesharwani et al., 2025b; Liao et al., 2019). ***Supplementary File 1*** contains comprehensive methodologies and procedures for transcriptome research.

### Identification of unique gene IDs and subsequent Gene Ontology (GO) analysis

DEGs in irradiated mice’s hippocampus were identified with a FDR cut-off of 0.05 (p values < 0.05) and a log2FC greater than 1.0. GO analysis was carried out using the Database for Annotation, Visualization, and Integrated Discovery (DAVID) version 2024q2 to identify important genes and pathways involve with irradiation-associated neurotoxicity, also the target gene for DH (Mi et al., 2017). Pathway enrichment analysis was performed on DEGs and DEPs affected by irradiated and/or DH therapy using the DAVID bioinformatics tool (p < 0.05) (Culhane et al., 2005; Kesharwani et al., 2025b; Mi et al., 2017; Singh et al., 2025).

### Proteomics study

The untargeted proteome was examined using Nano Liquid Chromatography (Q-Orbitrap) Mass Spectrometry (LCHRMS). Animals were sacrificed using cold saline, and their brains were rapidly collected. The hippocampus was isolated and protein extracted using a lysis solution containing 4% sodium dodecyl sulphate, 100 mM Tris-HCl (pH 8.5), 10 mM tris (2-carboxyethyl) phosphine, 40 mM 2-chloroacetamide, and a HALT protease/phosphatase inhibitor cocktail. Extracted protein were digested normally with trypsin in 55 mM acetic acid (1:50 ratio). The samples were subsequently vacuum dried in the vacuum concentrator for an hour and a half at 30°C in the AH-L mode. The collected samples were maintained at –80°C prior to LC-MS analysis (Kesharwani et al., 2025b; Prasad et al., 2022; Singh et al., 2025; van Oostrum et al., 2023). Detailed protocols for proteome analysis are provided in ***Supplementary File 1*.**

### ELISA Assay

The levels of 6-phosphofructokinase (PFKM: *MyBiosource, MBS099507*), glucose-6-phosphate dehydrogenase (G6PDH: *Sigma, MAK015*), hexokinase (HK), pyruvate dehydrogenase (PDH: *MyBiosource, MBS9719209*), lactate dehydrogenase (LDH: *MyBiosource, MBS034493),* α-ketoglutarate dehydrogenase (α-KGDH: *ImmunoTag, ITBC0083*), pyruvate: (*ImmunoTag, ITFA1016*) and lactate: (*ImmunoTag, ITFA1177*) in tissue homogenates of hippocampus were quantified using a following the manufacturer’s instructions. Detailed methodologies are given in ***supplementary 1***.

### Immunostaining, imaging, and confocal microscopic analysis

After saline perfusion, mice were transcardially perfused with 4% paraformaldehyde (PFA) in 0.1 M phosphate buffer (pH 7.4). The brains were carefully isolated from the skull and post-fixed in 4% PFA overnight at 4°C. Subsequently, the brains were dehydrated in a graded sucrose solution (10-30% w/v) until they sank, and sectioned into 30-μm-thick slices. The sections were immunostained to evaluate neuronal glucose transporters (MAP-2: *Abcam, ab254264* and GLUT3: *G-Biosciences, ITT05463*) and microglial glucose transporters (IBA-1: *G-Biosciences, ITT06442* and GLUT5: *G-Biosciences, ITT13545*) in the hippocampus. For each sample, Z-stacks of three immunostained coronal slices were captured using a Carl Zeiss LSM 880 confocal microscope at ICMR-RMRIMS, Patna. Comprehensive protocols for immunohistochemistry and confocal microscopy are provided in ***Supplementary File 1*** (Kesharwani et al., 2025a; Kesharwani et al., 2025b; Kesharwani, 2026; Parihar et al., 2015; Parihar and Limoli, 2013; Singh et al., 2025).

### Assessment of neurotransmitter content

The hippocampus was homogenized in 400□µL methanol with 100□ng/mL isoprenaline, followed by centrifugation at 12,000□rpm for 10□minutes. After vacuum drying the supernatant for 2□hours, the sample was reconstituted in 200□µL methanol, mixed well, and centrifuged at 12,000□rpm for 5□minutes. A 100 µL portion of the supernatant was transferred to sample vials for the analysis of glutamate, glutamine, GABA, dopamine, and kynurenic acid using the LC-HRMS system (He et al., 2013; Kesharwani et al., 2025a; Kesharwani et al., 2025b; Kesharwani, 2026; Lim et al., 2023). Further details on neurotransmitter quantification are provided in ***Supplementary File 1***.

### Data Analysis and Statistics

Graphs and figures were prepared using GraphPad Prism 8.0.1. Various statistical analyses were applied to different experiments, as described in the respective sections. Transcriptomics and proteomics data were processed using group comparisons, made against sham-irradiated mice, and differentially expressed genes and proteins were considered significant at an false discovery rate (FDR) of 0.05 (indicating a 95% probability of differential expression) and an absolute log2 fold-change. In Proteome Discoverer, differential expression p-values were calculated to assess statistical significance using a T-test, followed by adjustment with the Benjamini-Hochberg method. Q-values for differential expression were determined using scores from the decoy database search. For immunohistochemistry, neurotransmitter, and behavioral data, statistical analysis was performed using a two-way ANOVA with Tukey’s multiple comparisons test. with irradiation and DH as the independent variables. Data are presented as mean ± standard error of the mean (SEM). Statistical significance was defined as *p < 0.05, **p < 0.01, ***p < 0.001, and **p < 0.0001.

## Results

### DH treatment attenuates radiation-associated learning and memory impairments in NOR, OiP, and TO tests

Analysis using two-way ANOVA of the total time spent exploring both objects in the NOR test showed no significant interaction among irradiation and DH (F_1, 36_ = 1.786, p = 0.190) also no main impact of irradiation (F_1, 36_ = 0.023, p = 0.8794) and DH (F_1, 36_ = 3.839, p = 0.0579). This suggests that irradiation and/or DH treatment did not affect locomotor activity. (**Fig. 2A1**). Three-way ANOVA analysis demonstrated a significant triple interaction between novelty preference, irradiation, and DH treatment (F□,□□= 30.03, p < 0.0001) and significant two-way interactions between novelty preference and irradiation (F□,□□= 69.83, p < 0.0001) and novelty preference and DH (F□,□□= 21.67, p < 0.0001). Both unirradiated groups (0 Gy ± DH) exhibited a clear preference for novelty, as indicated by the significantly greater percentage of time spent exploring the novel object (0 Gy + Veh, p < 0.0001; 0 Gy + DH, p < 0.0001; **Fig. 2A2**). In contrast, irradiated mice receiving vehicle showed no preference for novelty (p = 0.981; **Fig. 2A2**), consistent with previous reports demonstrating that cranial irradiation significantly impairs object recognition memory (Acharya et al., 2016). Importantly, irradiated mice treated with DH retained the ability to distinguish the novel object, showing a significant preference for novelty (p < 0.0001; **Fig. 2A2**), indicating that DH mitigates radiation-induced memory deficits. Two-way ANOVA of the DI revealed a significant interaction between irradiation and DH (F□,□□= 33.760, p < 0.0001), along with significant main effects of irradiation (F□,□□= 6.072, p = 0.0186) and DH treatment (F□,□□= 23.16, p < 0.0001; **Fig. 2A3**). Tukey’s post hoc test showed that DH-treated irradiated mice displayed significantly higher DI compared with vehicle-treated irradiated mice (p < 0.0001; **Fig. 2A3**), indicating preserved memory function.

**Fig. 2:**
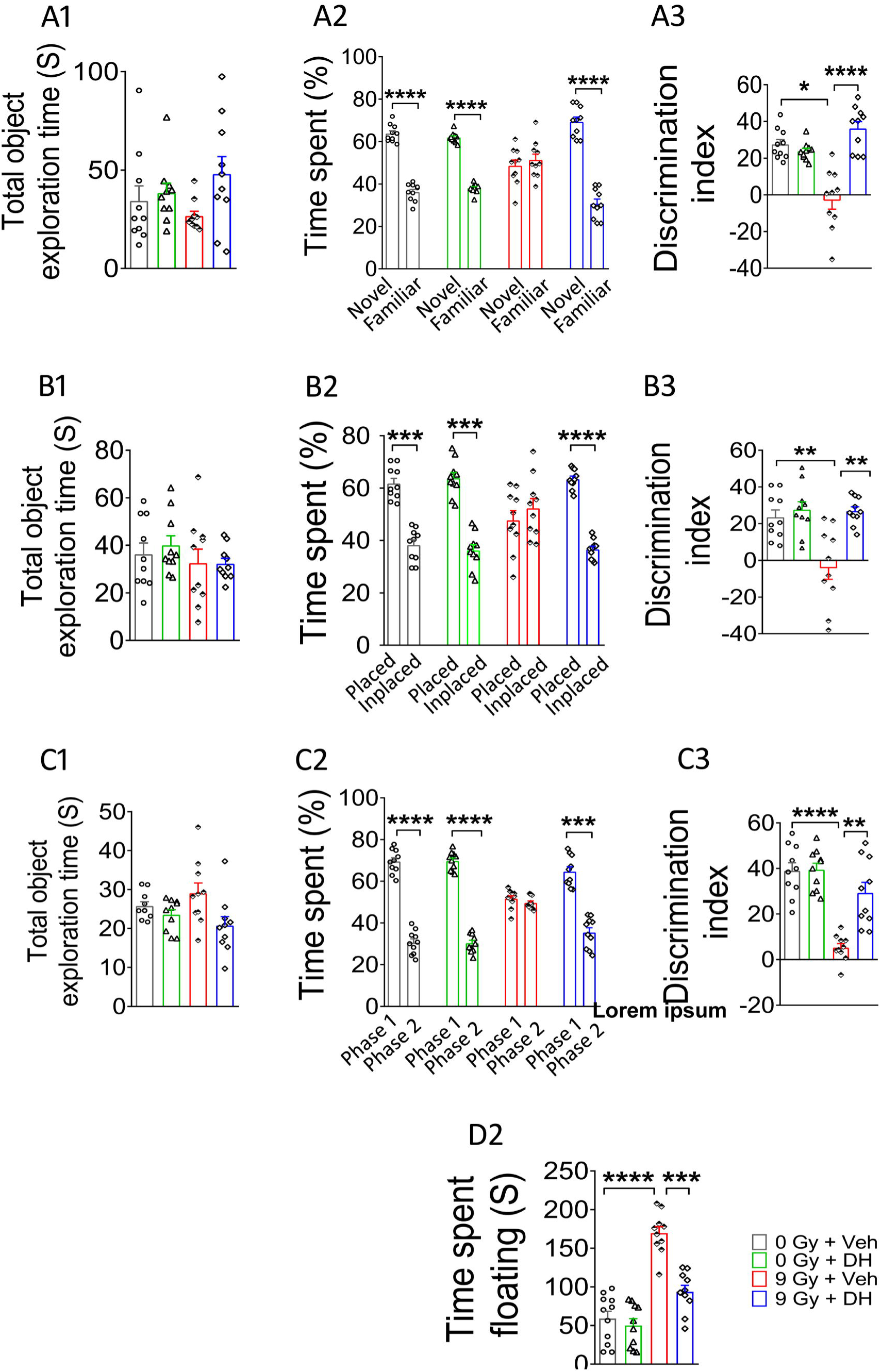
DH mitigates irradiation-induced cognitive deficits. Total exploration time remained unchanged across groups in NOR (A1), OiP (81), and TO (C1). Irradiation reduced novelty preference, as shown by decreased discrimination index (DI) and exploration of novel/relocated objects in NOR (A2-A3), OiP (82-83), and TO (C2-C3); these deficits were reversed by DH treatment. In the FST, irradiation increased floating time (D1), indicating depression-like behavior, which was significantly reduced by DH (D2). Data were analyzed by two-way or three-way ANOVA with Tukey’s post hoc test. *p < 0.05, **p < 0.01, ***p < 0.001, ****p < 0.0001; n = 10/group.

The OiP task, which is recognised as likewise depend on intact hippocampus and perirhinal cortex-dependent brain function, was administered to acclimated mice after the NOR task. All three factors-irradiation’s main effect (F_1, 36_ = 0.159, p = 0.693), DH’s effect (F_1, 36_ = 0.15, p = 0.693), and the interaction between the two variables were not statistically significant when the total time spent examining the two items was analysed using two-way ANOVA (F_1, 36_ = 0.200, p = 0.657) (**Fig. 2B1**). This suggests that exposure to irradiation and/or DH did not hinder the natural exploratory behavior of mice during the OiP task. Three-way ANOVA of the time spent exploring familiar and novel object locations revealed a significant triple interaction among novelty preference, irradiation, and DH treatment (F□,□□= 15.82, p = 0.0002), as well as significant two-way interactions between novelty preference and irradiation (F□,□□= 26.78, p < 0.0001) and between novelty preference and DH treatment (F□,□□= 17.68, p < 0.0001). Post hoc analysis using Tukey’s multiple comparisons test showed that both unirradiated groups (0 Gy ± DH) displayed a clear preference for objects moved to novel locations, indicated by a greater percentage of time spent exploring the new locations. In contrast, irradiated vehicle-treated mice showed no such preference. Importantly, irradiated mice treated with DH retained the ability to distinguish novel object locations (0 Gy + Veh, p < 0.0001; 0 Gy + DH, p < 0.0001; 9 Gy + Veh, p = 0.883; 9 Gy + DH, p < 0.0001; **Fig. 2B2**). For the discrimination index (DI), two-way ANOVA revealed significant main effects of irradiation (F□,□ □= 8.23, p = 0.0068) and DH treatment (F□,□ □= 14.20, p = 0.0006), as well as a significant interaction between irradiation and DH (F□,□ □= 8.68, p = 0.0056). Tukey’s multiple comparisons test demonstrated that DH significantly mitigated radiation-induced deficits, as evidenced by higher DI in irradiated mice treated with DH compared with vehicle-treated irradiated mice (p = 0.0002; **Fig. 2B3**).

Subsequent to the OiP task, we engaged in the TO task, which is recognized for its dependence on the integrity of the perirhinal cortex and functions reliant on the entorhinal cortex. There was no significant interaction or main effect of radiation (F_1, 36_ = 0.0149, p =0.9035) or DH (F_1, 36_ = 7.457, p = 0.0097) when analysing the total time spent examining both objects in the test phase using two-way ANOVA (**Fig. 2C1**). Similar to previous task, these data indicate that exposure to radiation and/or DH treatment did not impair locomotor activity on the TO task. Three-way ANOVA of the time spent exploring familiar and novel objects revealed a significant triple interaction among novelty preference, radiation, and DH treatment (F□,□□= 24.18, p < 0.0001), as well as significant two-way interactions between novelty preference and radiation (F□,□□= 84.65, p < 0.0001) and between novelty preference and DH treatment (F□,□□= 26.46, p < 0.0001). Post hoc analysis using Tukey’s multiple comparisons test showed that both unirradiated groups (0 Gy ± DH) displayed a clear preference for the novel object, indicated by a greater percentage of time spent exploring it (0 Gy + Veh, p < 0.0001; 0 Gy + DH, p < 0.0001; **Fig. 2C2**). In contrast, irradiated mice receiving vehicle showed no preference for novelty (9 Gy + Veh, p = 0.377; Fig. 2C2). Importantly, irradiated mice treated with DH retained the ability to discriminate the novel object (9 Gy + DH, p < 0.0001; **Fig. 2C2**), indicating that DH mitigates radiation-induced deficits in object recognition. For the discrimination index (DI), two-way ANOVA revealed significant main effects of radiation (F□,□□= 42.32, p < 0.0001) and DH treatment (F□,□□= 13.23, p = 0.0009), as well as a significant interaction between radiation and DH (F□,□□= 12.09, p = 0.001). Tukey’s multiple comparisons test indicated that DH significantly alleviated the diabetes-associated recency deficit, as evidenced by a higher DI in DH-treated irradiated mice compared with vehicle-treated diabetic mice (p < 0.0001; **Fig. 2C3**).

Additionally, the immobility time (the amount of time the mouse spent floating) was used to measure depressive-like behavior in the FST. Two-way ANOVA on floating time showed that there was a potential interaction among DH and HFD-STZ F_1, 36_ = 15.72, p = 0.0003), as well as a significant effect of irradiation (F_1, 36_ = 79.10, p < 0.0001), and DH F_1, 36_ = 21.16, p < 0.0001). The results of Tukey’s multiple comparison revealed that the irradiated mice showed enhanced depressive-like behavior, as evidenced by a potential increase in floating time (p < 0.0001, 0 Gy + Veh vs. 9 Gy + Veh). However, irradiated mice who received DH treatment showed a decrease in depressive-like behavior. This was evident from the significantly reduced floating time observed after DH treatment, compared to the group that received a unirradiated mice without DH treatment (p = 0.0002, **Fig. 2D2**).

### Radiated mice showed an improvement in spatial learning and memory after receiving DH therapy

All groups of mice underwent the fixed platform Water Maze Test (WMT) to evaluate their spatial learning and memory retrieval capabilities. A two-way analysis of variance showed that there was no difference in swimming speeds across groups that were treated with radiation or DH (p = 0.9701; **Fig. 3A2**). Further, for the latency to escape platform, a two-way ANOVA determined a potential day’s effect (F_6, 252_ = 514.1, p < 0.0001), group effect (F_3, 252_ = 187.9, p < 0.0001), and significant interaction between days and group (F_18, 252_ = 14.93, p < 0.0001).Further, Tukey’s multiple comparison test revealed that irradiation significantly alters the spatial learning (p < 0.0001), as indicated by increased latencies to reach the platform, whereas DH treatment significantly enhanced the spatial learning in irradiated mice, as indicated by decreased latencies to reach the platform (p < 0.0001,**Fig. 3A3).**

**Fig. 3:**
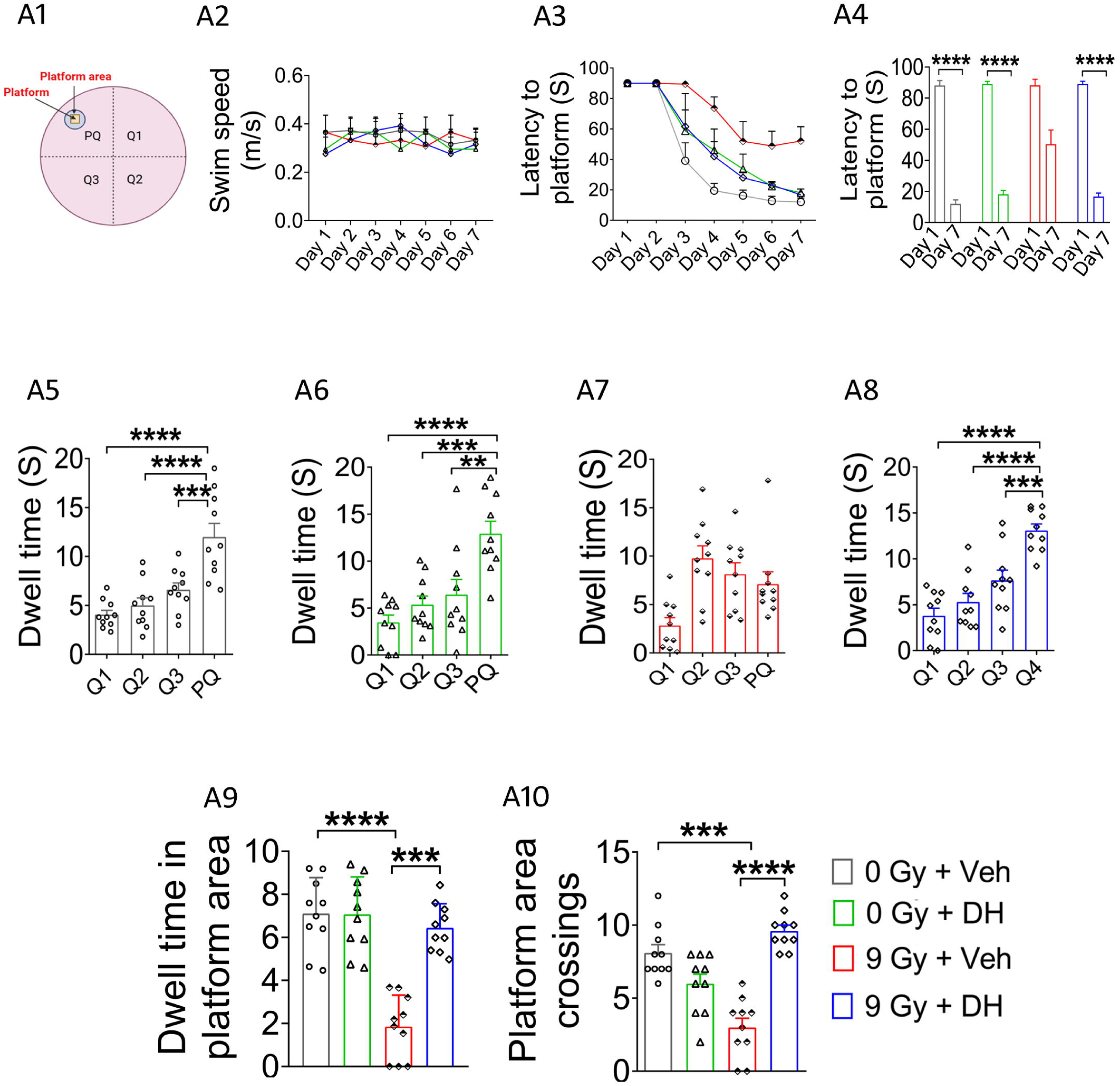
DH treatment promotes spatial learning and memory in cranial irradiated mice: Schematic diagram of the pool, platform and quadrants (A1). There were no differences between the groups for swim speed (A2). Latency to escape platform shows the lesser affinity of irradiated mice towards the platform quadrant, and DH treatment potentially improved it (A3-A4). Probe test conducted 24 hours after the last learning session revealed similar results (A5-A8). Irradiated group (A?) showed lesser affinity for the platform quadrant, in comparison to the other groups, while DH treated group (AS) showed greater affinity toward the platform quadrant, in comparison to other three quadrants of pool. Additional parameters of memory retrieval ability such as the dwell time in the platform area (A9), and platform area crossings (A10) were also comparable between each group. PQ (NE-Q), platform quadrant (northeast quadrant); SE-Q, southeast quadrant; SW-Q, southwest quadrant; and NW-Q, northwest quadrant. *p < 0.05, **p < 0.01; ***p < 0.001.

A direct comparison of latency to reach the platform between training session day 1 and day 7, unpaired t-test revealed that irradiation potentially impaired the performance of the irradiated mice (p = 0.0166), while DH treatment led to a notable improvement (p < 0.0001; **Fig. 3A4).** Data analysis from the memory retrieval assessment conducted 24 hours post the final learning session revealed a diminished capacity in irradiated mice, while DH-treated mice exhibited maintained proficiency in spatial memory formation. One-way Anova suggested irradiated mice treated with vehicle showed no affinity for PQ over other quadrants (p = 0.7090), while DH-treated animals showed a significant preference for the PQ over all other quadrants (p < 0.0001) **Fig. 3A6 and A8**).

Furthermore, for the time spent in platform, two way ANOVA suggested a significant impact of irradiation (F_1, 36_ = 26.55, p < 0.0001), DH effect (F_1, 36_ = 29.58, p < 0.0001), and a potential interaction among irradiation and DH (F_1, 36_ = 15.06, p = 0.0004).Tukey’s multiple comparison test showed that irradiation potentially impaired the performance of irradiated mice (p < 0.0001) as evidenced by reduced time spent in platform, while DH-treated irradiated group allocated more time within the platform quadrant compared to the irradiated groups (p < 0.0001; **Fig. 3A9**).

### Analysis of differentially expressed transcripts (DEGs)

To examine whether energy metabolism in irradiated mice is associated with DEGs, we performed transcriptome analysis of the hippocampus by RNA sequencing approach. Transcriptomic studies identified consisting of 318 differentially expressed genes, among which 179 genes were differentially expressed between irradiated mice vs control (9 Gy + Veh vs 0 Gy + Veh) and 139 genes were differentially expressed in irradiated mice vs DH treated mice (9 Gy + DH vs 9 Gy + DH) (***Supplementary File 2***). Genes exhibiting an absolute log□fold-change ≥1 with p < 0.05 were considered differentially expressed, with positive values classified as upregulated and negative values as downregulated (**Fig. 4A1**). According to our RNA-seq data, 17 genes were either upregulated (112genes) or downregulated (67 genes) in irradiated mice compared with control mice (9 Gy + Veh vs 0 Gy + Veh) (**Fig. 4A1**). Similarly, 23 genes were upregulated and 111 genes were downregulated in irradiated mice when compared with irradiated mice received DH treatment (9 Gy + DH vs 9 Gy + Veh) (**Fig. 4A2**). Gene Ontology analysis revealed that the majority of the DEGs encoded genes related to immune responses, immunity, ATP synthesis, energy metabolism, mast cell activation, mitochondrial functions *(Bc1, Igha, Cbx3-ps4, Cbx3-ps2, Phxr4, Cd6, mt-Td, Cox6c2).* Furthermore, GO analysis revealed significant changes in the clusters of genes associated with neuropeptide receptor activity, cholesterol metabolism, cell migration, inflammation, GABA receptor, neurodevelopmental disorder, cell proliferation and functions and lipid metabolism (*Gpr165, Idi1-ps1, Nr4a3, Lbp, Gabrb2, Pcdhgb1, Prkcd and Plin4*). Moreover, DH treatment significantly regulated hallmark genes (*mt-Nd4l, mt-Nd5, mt-Atp8, mt-Td, Pin4-ps, Atp5pb-ps, mt-Nd1, mt-Atp6, mt-Nd2, Gbp11, Oacyl, Prg4 and Cd180*) associated with mitochondrial respiratory chain, mitochondrial ATP functions, neurotransmission function, inflammation, neural differentiation and learning and memory (**Fig. 5**). In addition, 177 DEGs related to irradiated mice were mapped and list of significant pathways largely involved in biological processes, cellular components, molecular functions, translational modifications, and KEGG pathways ATP metabolic process, negative regulation of neuron apoptotic process, ion transport, neuron projections, ion channel activity, Aminoacyl-tRNA biosynthesis. Further, DH treatment influenced numerous pathways in irradiated mice, with the greatest effect on neuroinflammation, immunity, energy metabolism, mitochondrion and pathways of neurodegeneration impacted all the foregoing pathways (**Fig. 5**). The details of the individual significance values for the enriched KEGG pathways are provided in ***Supplementary File 3*.**

**Fig. 4:**
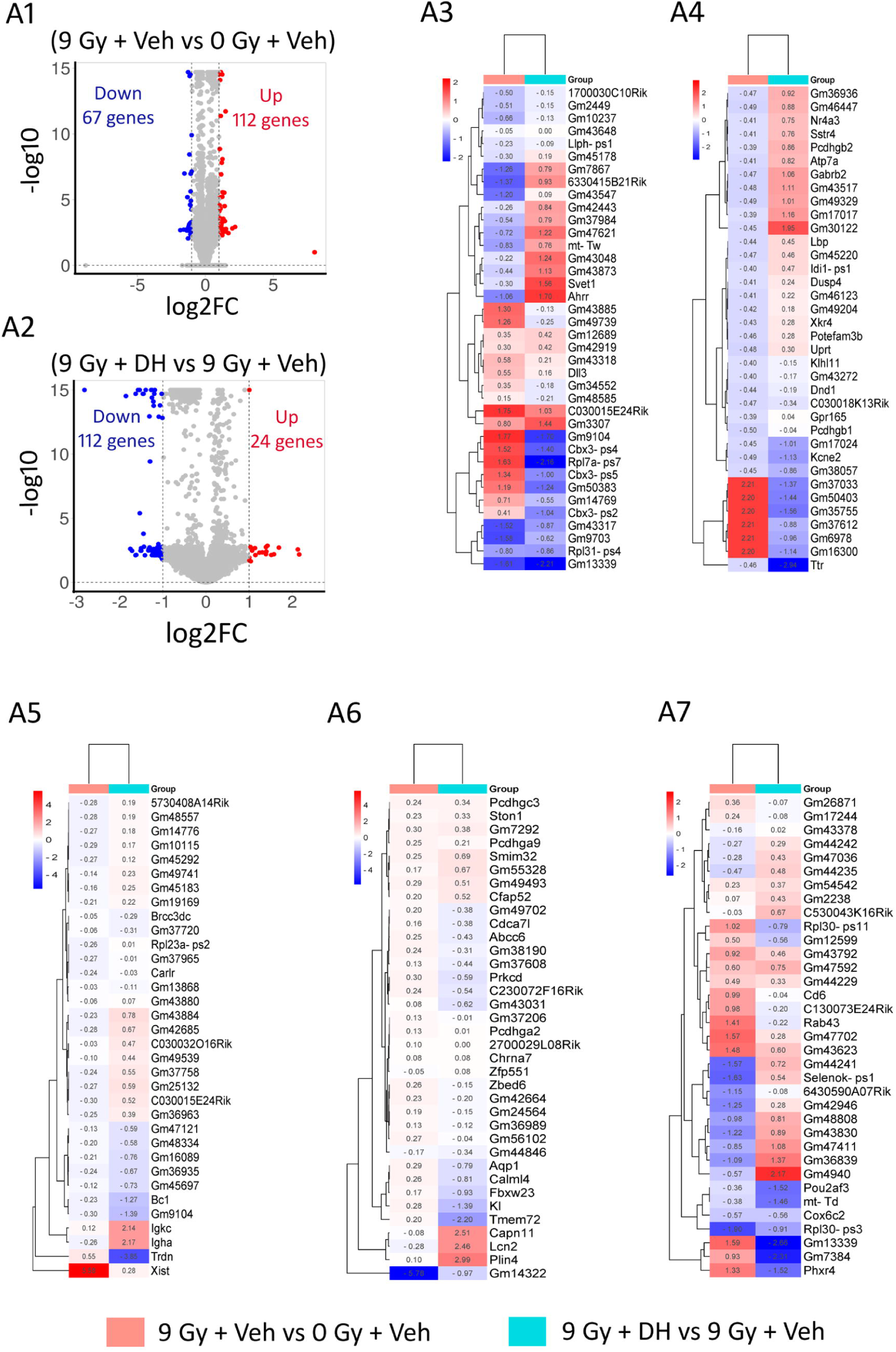
Volcano plots for comparative analysis of unirradited mice and irradiated mice (0 Gy + Veh vs 9 Gy + Veh) (A1), irradiated mice with vehicle or irradiated mice with DH (9 Gy + veh vs 9 Gy + DH) (A2). Colored symbols represent differentially expressed genes (DEGs) (Padj <0.05): Red, upregulated; and blue, downregulated. Heatmaps showing the effects of irradiation and DH treatment, as comparative analysis between the 9 Gy + Veh vs 0 Gy + Veh and 9 Gy + DH vs 9 Gy + Veh. Numbers represent the log2 value for each gene (A3-A7) **N=** 3 per group.

**Fig. 5:**
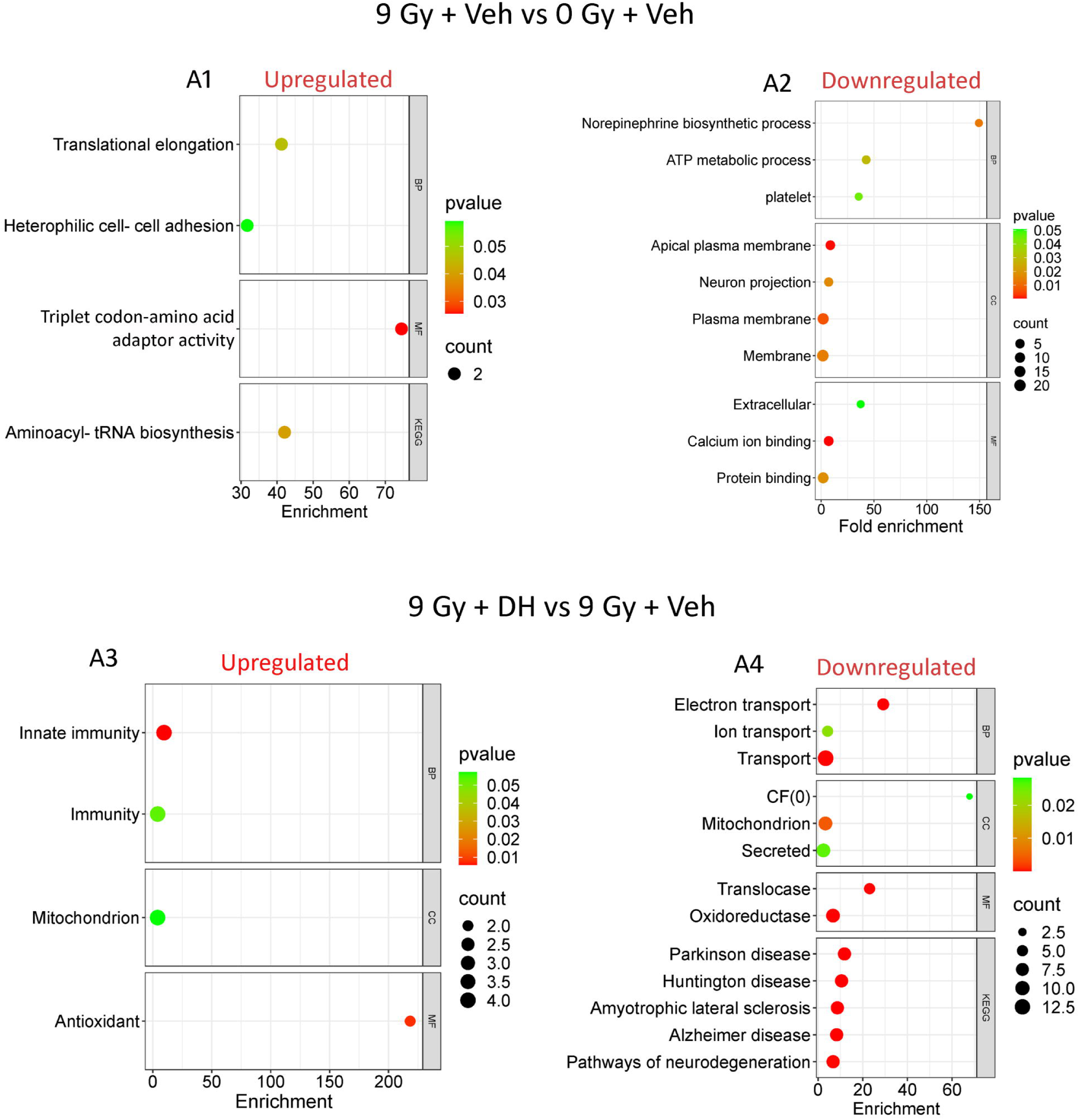
Irradiated mice display altered a pathway of brain. GO categories showing enrichment of DEGs that were upregulated (A1) or down-regulated (A2), in irradiated mice from comparative analysis with unirradiated (9 Gy + Veh vs O Gy + Veh), while the DH regulates the altered pathways, showing enrichment of DEGs that were upregulated (A3) or down-regulated (A4), in DH treated mice from comparative analysis with irradiated (9 Gy + Veh vs 9 Gy + Veh) **N=** 3 per group.

**Fig. 6:**
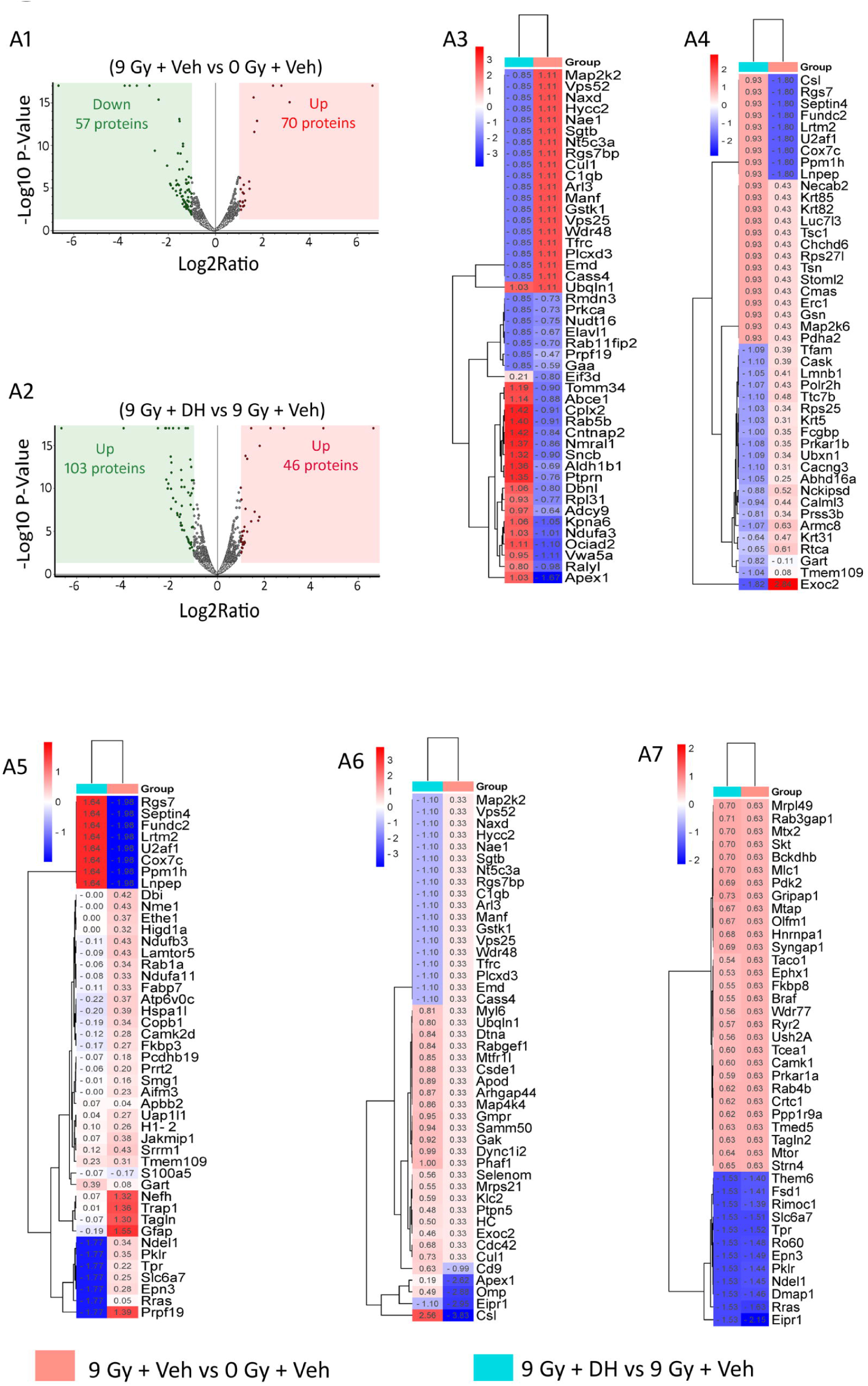
Volcano plots for comparative analysis of between irradiated and unirradiated (mice 0 Gy + Veh vs 9 Gy + Veh) (A1), irradiated mice with vehicle or irradiated mice with DH (9 Gy + DH vs 9 Gy + Veh) (A2). Colored symbols represent differentially expressed proteins (DEPs) (Padj <0.05): Red, upregulated; and blue, downregulated. Heatmaps showing the effects of irradiation and DH treatment, as comparative analysis between the 9 Gy + Veh vs 0 Gy + Veh and 9 Gy + DH vs 9 Gy + Veh. Numbers represent the log2 value for each gene (A3-A7). N= 3 per group.

### Differential protein expression analysis (DEPs)

Proteomic analysis uncovered 2172 proteins, out of which 128 showing differential expression in irradiated mice comparison to controls (9 Gy + Veh vs 0 Gy + Veh), or 150 differential expression irradiated mice treated with DH (9 Gy + Veh vs 9 Gy + DH) ***(Supplementary File 4 and* *Fig. 5A1 and* *5B1)***. The DEPs identified in irradiated mice exhibit several characteristics commonly related to neurotoxicity, like mitochondrial dysfunction, energy metabolism, and neurotransmitter regulation. These factors play a key role in the development and advancement of neurodegenerative diseases, as shown in Figure 5C1 and 5C2 (*Syngap1, Pdk2, Mtx2, Apod, Prkar1a, Taco1, Map2k2, Gfap*). The functional enrichment analyses revealed that the DEPs in irradiated mice were found to play roles in various biological processes and pathways. These processes and pathways include ATP metabolic process, neuronal projection, mitochondria activity, ATP binding, cell projection, cytoplasm integrity, synaptic function, metabolic processes, immune system processes, and inflammatory signalling **(Fig. 7)**. Treatment significantly affected various pathways in irradiated mice, with prominent effects on metabolic and energy-related processes. The enriched KEGG pathways detailed individual significance values can be found in **Supplementary File 5**.

**Fig 7:**
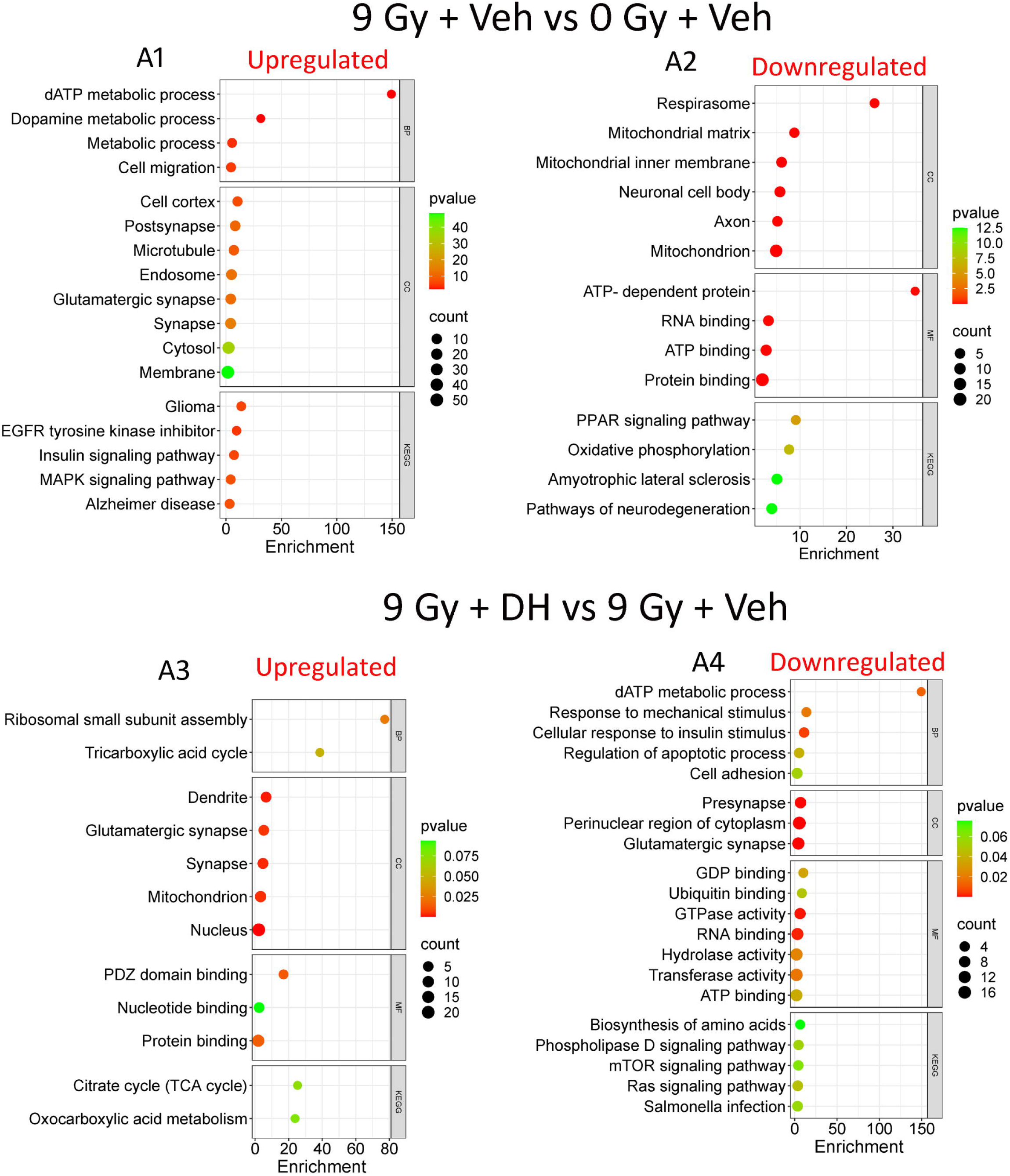
Irradiated mice display altered a pathway of brain. GO categories showing enrichment of DEPs that were upregulated (A1) or down-regulated (A2), in irradiated mice from comparative analysis with unirradiated (9 Gy + Veh vs O Gy + Veh), while the DH regulates the altered pathways, showing enrichment of DEPs that were upregulated (A3) or down-regulated (A4), in DH treated mice from comparative analysis

### Irradiation disrupts glucose metabolism and energy production in the hippocampus by affecting glycolytic enzymes including 6-phosphofructokinase (PFKM), glucose-6-phosphate dehydrogenase (G6PDH) and hexokinase (HK)

An ELISA assay was performed to assess the impact of radiation on glycolytic enzymes activity. Two-way ANOVA of PFKM expression indicated a significant main impact of irradiation (F_1, 16_ = 10.10, p = 0.006) and DH (F_1, 16_ = 52.37, p < 0.0001), also a potential interaction among irradiation and DH (F_1, 16_ = 7.673, p = 0.014). Additionally, Tukey’s multiple comparison indicated a potential decrease in PFKM activity (p < 0.0001) in the prefrontal cortex of irradiated mice, whereas treatment with DH led to a notable increase in PFKM activity (p = 0.0034) in the same group of irradiated mice ***(Fig. 8A1)***. Further, irradiation potentially altered G6PDH activity in irradiated mice brain. A two-way ANOVA performed on G6PDH activity indicated a potential interaction among irradiation and DH (F_1, 16_ = 29.72, p < 0.0001), a main effect of irradiation (F_1, 16_ = 22.32, p = 0.0002), and a significant effect of DH (F_1, 16_ = 12.49, p = 0.0028). Additionally, Tukey’s multiple comparison test indicated a possible decrease in G6PDH activity (p < 0.0001) in the irradiated brain, while DH treatment may have the potential to restore its activity (p < 0.0001) in the brains of irradiated mice ***(Fig. 8A2)***. In contrast, this irradiation did not influence HK activity in the brains of mice. Two-way ANOVA assay of HK indicated no potential interaction between irradiation and DH (F_1, 16_ = 1.126, p = 0.3043), main effect irradiation (F_1, 16_ = 0.1646, p = 0.6903) and DH (F_1, 16_ = 0.6352, p = 0.4371, ***Fig. 8A3)***.

**Fig 8:**
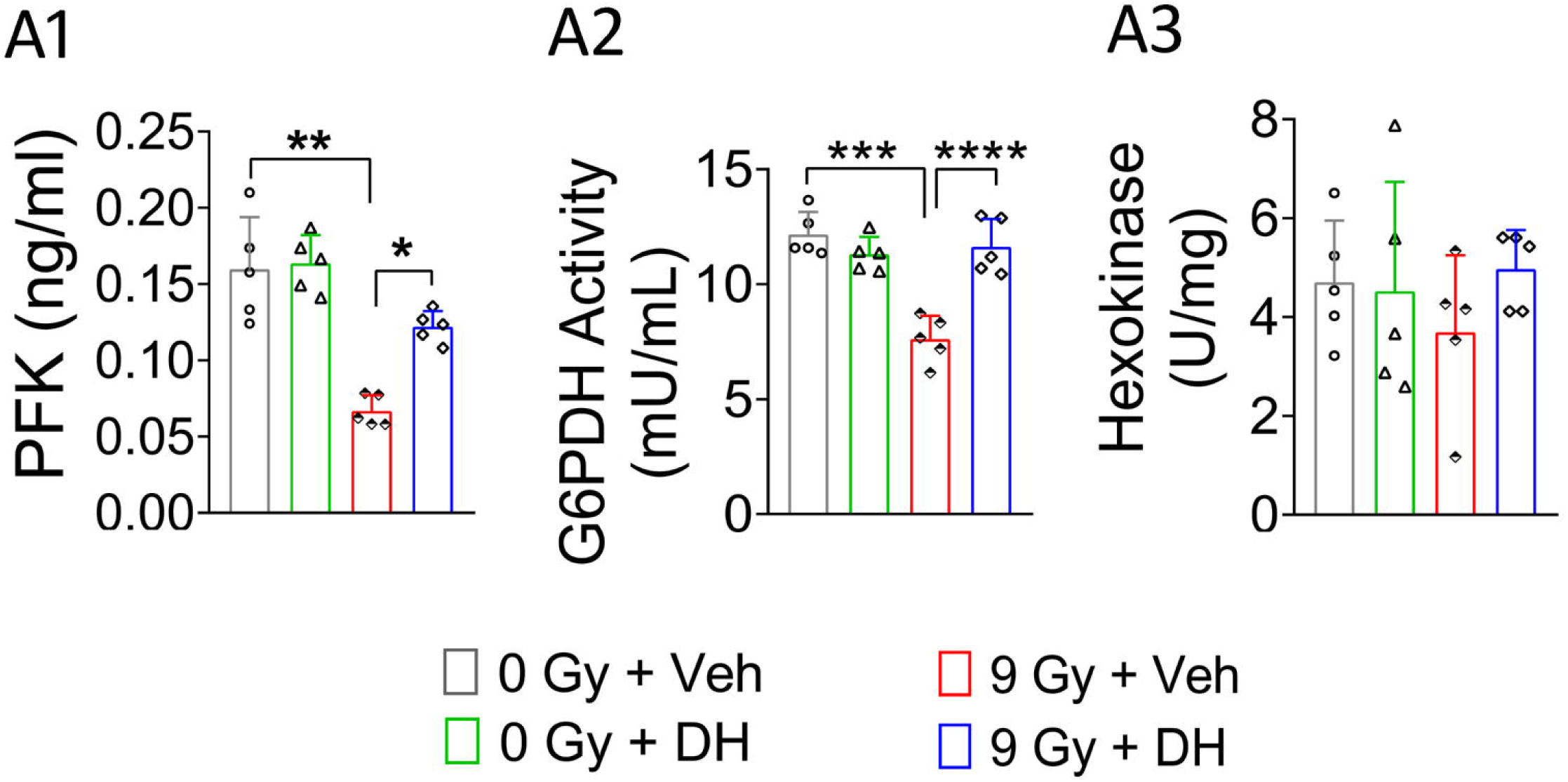
Irradiation significantly reduced the activity of PFKM and G6PDH in the brain, while DH treatment partially restored their activity (A1-A2). In contrast, HK activity remained unchanged following irradiation (A3). Data are presented as mean ± SEM (n = 5 per group). Statistical analysis was performed using two-way ANOVA followed by Tukey’s multiple comparisons test ***p < 0.001, ****p < 0.0001.Two-way ANOVA followed by Tukey’s multiple comparison. ***p < 0.001, ****p < 0.0001. **N** = 5 per

### Irradiation alters activity of enzymes regulating mitochondrial energetic and energy metabolism in brain

To evaluate the impact of irradiation on energy metabolism and ATP synthesis, the essential mitochondrial enzymes (pyruvate dehydrogenase (PDH), lactate dehydrogenase (LDH), and alpha-ketoglutarate dehydrogenase (α-KGDH) and substrates were analysed by ELISA. The Two-way ANOVA of PDH activity assay revealed a potential effect of irradiation (F_1, 16_ = 5.147, P = 0.0375) and DH (F_1, 16_ = 26.27, p = 0.0001), also significant interaction among irradiation and DH (F_1, 16_ = 57.03, p < 0.0001). Further, Tukey’s multiple comparison showed reduced activity of PDH in the PFC of irradiated brain (p < 0.0001), while DH significantly increased its activity in irradiated mice (p < 0.0001, ***Fig. 9A1*)**. Altered PDH activity, resultant to its disturbed pyruvate level in hippocampus. Two-way ANOVA of pyruvate level indicated a potential impact of irradiation (F_1, 16_ = 14.53, p = 0.0015) and DH (F_1, 16_ = 10.65, p = 0.0049), as well as potential interaction among irradiation and DH (F_1, 16_ = 5.077, p = 0.0386). Tukey’s multiple comparison showed increased level of pyruvate in the PFC of irradiated brain (p = 0.0028), while DH potentially reduced its level in irradiated mice (p = 0.0063, ***Fig. 9A2*)**. Further, Two-way ANOVA of LDH activity in PFC of irradiated mice brain. Two-way ANOVA of LDH activity revealed a potential interaction (F_1, 16_ = 14.55, p = 0.0015) and main effect of irradiation (F_1, 16_ = 23.43, p = 0.0002) and DH (F_1, 16_ = 5.602, p = 0.0309). Further, Tukey’s multiple comparison revealed significant reduced LDH activity (p < 0.0001) in the PFC of irradiated mice while treatment with DH significantly increased LDH activity (p = 0.0024) in irradiated mice, ***Fig. 9A3*)**. Disturbed LDH activity resultant increased level of lactate level in PFC. Two-way ANOVA of lactate level indicated that potential interaction between irradiation and DH (F_1, 16_ = 15.17, p = 0.0013), main effect of irradiation (F_1, 16_ = 5.052, p = 0.0390) and DH (F_1, 16_ = 15.30, p = 0.0012). Tukey’s multiple comparison revealed increased level of lactate (p = 0.0002) in PFC of irradiated mice while treatment with DH reduced (p = 0.0025) its level in PFC of irradiated mice, ***Fig. 9A4*)**. The two-way ANOVA analysis of α-KGDH activity suggested that significant effect of irradiation (F_1, 16_ = 22.68, p = 0.0002) and DH effect (F_1, 16_ = 5.995, p = 0.0263) and significant interaction between irradiation and DH (F_1, 16_ = 23.17, p = 0.0002). Tukey’s multiple comparison test revealed irradiation significantly decreased α-KGDH activity (p < 0.0001), while DH treatment increased (p = 0.0005) its level in hippocampus of irradiated brain, ***Fig. 9A5*)**. These findings indicate that one of the potential underlying benefits of DH treatment is the ability to modulate PDH activity and its metabolic product pyruvate level.

**Fig. 9:**
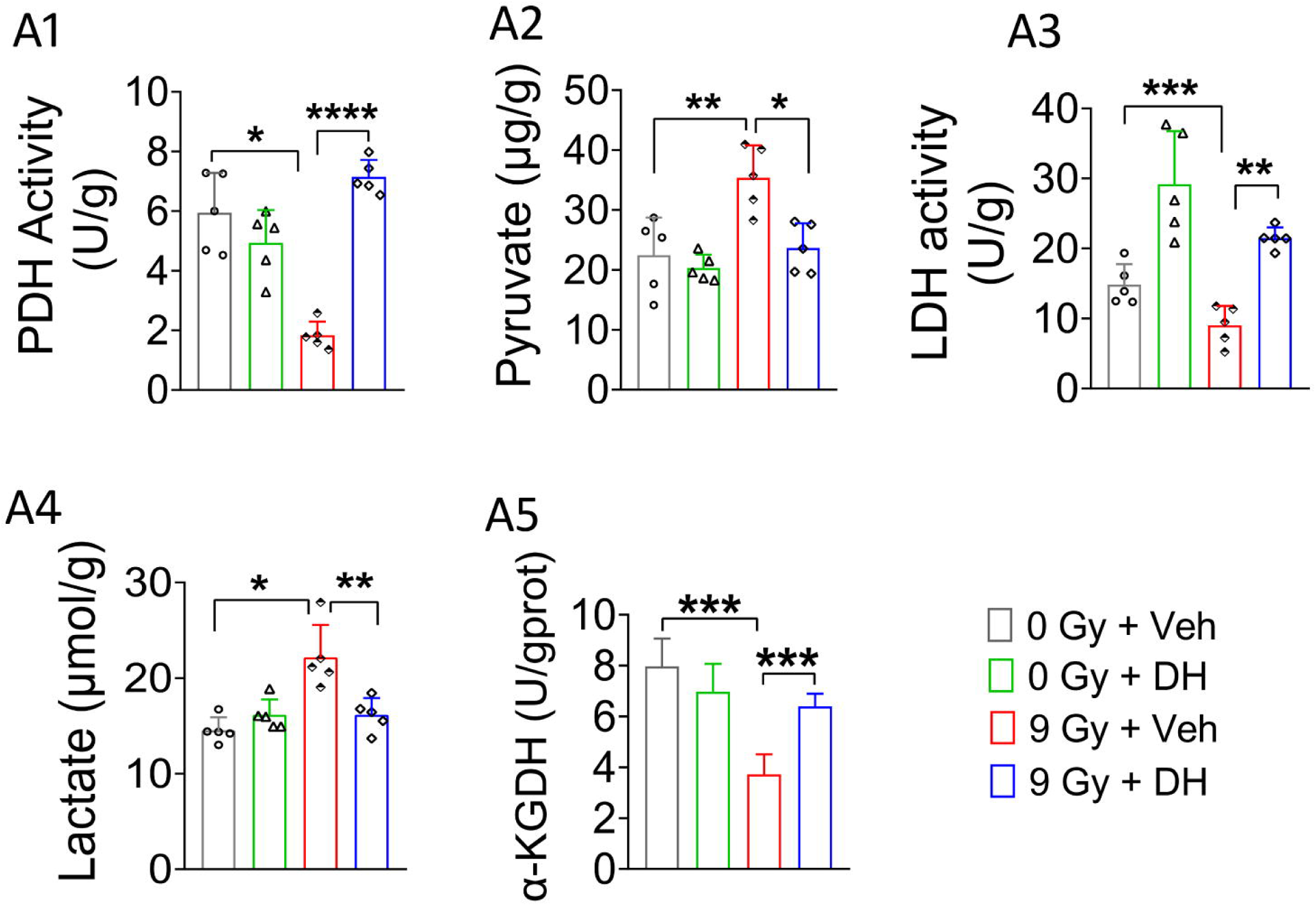
Irradiation disturbs activity of enzymes regulating mitochondrial energetic and energy metabolism in brain: Irradiated mice showed the reduced activity of PDH, LOH and a-KGDH, whereas DH treatment potentially regulated it (A1, A3 & AS). Disturbed activity of PDH and LOH resultant higher level of its metabolic product pyruvate and lactate in irradiated mice brain (A2 & A4). Two-way ANOVA followed by Tukey’s multiple comparison. ***p < 0.001,

### Irradiation potentially modulates GLUT5 expression in the hippocampus of irradiated mice

To ascertain impact of irradiation on Glucose transporters, we quantified the expression of GLUT5 expression in microglial cells (IBA1+) of the hippocampus of irradiated mice. Two-way ANOVA of GLUT5 showed a potential impact of irradiation (F_1, 12_ = 16.44, p = 0.0016) and DH (F_1, 12_ = 43.39, p < 0.0001), also a significant interaction among irradiation and DH (F_1, 12_ = 5.406, p = 0.0384). Tukey’s multiple comparison test showed reduced expression of GLUT5 in the hippocampus of irradiated mice (p = 0.0002), while DH treatment significantly promoted it (p = 0.0034). The findings imply that one possible mechanism underlying the benefits of DH treatment is the regulation of glucose uptake in microglial cells **(*Fig. 10*).**

**Fig. 10:**
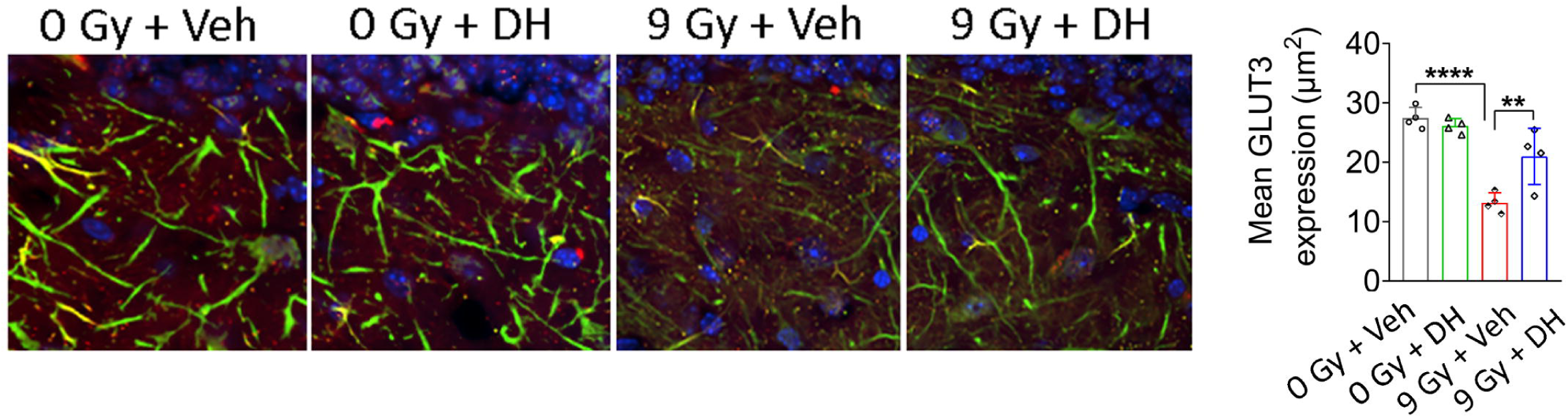
In irradiated mice, DH enhanced expression of neuronal glucose transporters and neuronal density in hippocampus. The quantification of MAP2 and GLUT3 + cells in the hippocampus of irradiated mice showed a significant reduction; however, DH treatment enhanced this severity (Green: MAP2+ neurons, Red; GLUT3+ glucose transporters, Blue: nuclear stain). Two-way ANOVA followed by Tukey’s multiple comparisons. *p < 0.05, **p < 0.01, ***p < 0.001, ****p< 0.0001. N = 4 per group.

### Irradiation significantly alters the expression of GLUT 3 in the hippocampus of irradiated mice

To ascertain impact of irradiation on Glucose transporters, we quantified the expression of GLUT3 in neuronal cells (MAP2 +) of the hippocampus of irradiated mice. Two-way ANOVA of GLUT3 expression revealed a significant effect of irradiation F_1, 12_ = 50.39, p < 0.0001) and DH (F_1, 12_ = 5.622, p = 0.0353), as well as a significant interaction between irradiation and DH F_1, 12_ = 11.14, p = 0.0059). Tukey’s multiple comparison test revealed lower expression of GLUT3 in the hippocampus of diabetic mice (p < 0.0001), while DH treatment significantly increased it (p = 0.0077). These data suggest that DH have potency to enhanced the expression of GLUT3 in hippocampus of irradiated mice **(*Fig. 11*)**.

**Fig. 11:**
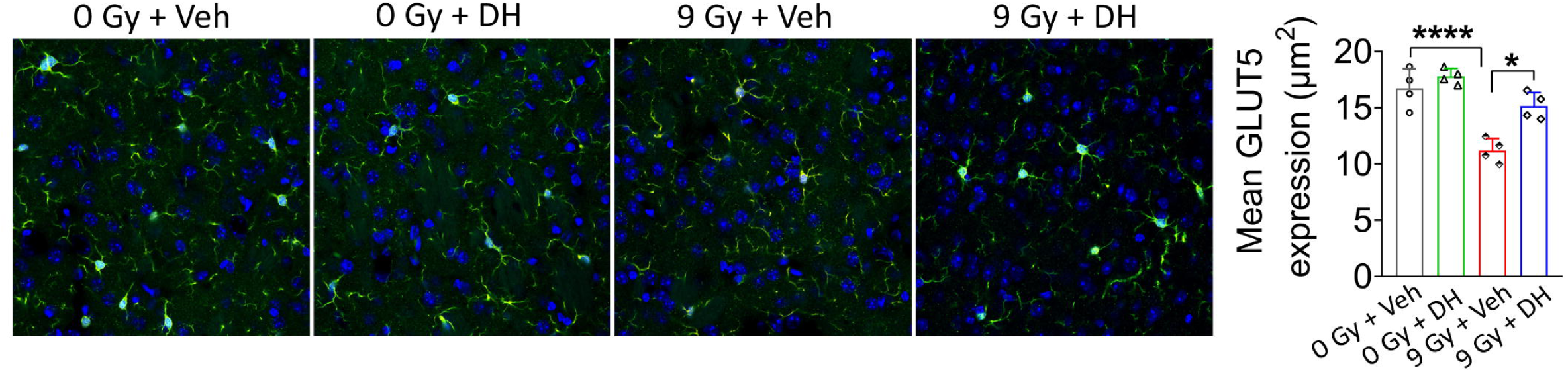
In irradiated mice, DH enhanced expression of microglial glucose transporters and no. of microglia in hippocampus. The quantification of IBA1 and GLUTS+ cells in the hippocampus of irradiated mice showed a significant reduction; however, DH treatment enhanced this severity (Green: IBA1+ microglia, Red; GLUTS+ glucose transporters, Blue: nuclear stain). Two-way ANOVA followed by Tukey’s multiple comparisons. *p < 0.05, **p < 0.01, ***p < 0.001, ****p< 0.0001. **N** = 4 per group.

### Irradiation alters the glutamine-glutamate/GABA cycles in the hippocampus

Two-way ANOVA of glutamine, glutamate, and GABA levels in the hippocampus revealed significant effect of irradiation (glutamine: F_1, 16_ = 7.269, p = 0.016; glutamate: F_1, 16_ = 43.85, p < 0.0001; GABA: F_1, 16_ = 8.527, p = 0.0100) and DH (glutamine: F_1, 16_ = 11.76, p = 0.003; glutamate: F_1, 16_ = 15.34, p = 0.0012; GABA: F_1, 16_ = 21.88, p = 0.0003), as well as significant interaction between irradiation and DH (glutamine: F_1, 16_ = 14.02, p = 0.0018; glutamate: F_1, 16_ = 4.546, p = 0.0488; GABA: F_1, 16_ = 25.49, p = 0.0001). Tukey’s multiple comparison revealed a decrease in glutamine and GABA in irradiated mice (glutamine: p = 0.0012 and GABA: p = 0.0002 for 0 Gy + Veh vs 9 Gy + Veh), while DH treatment increases (glutamine: p = 0.0006 and GABA: p < 0.0001 for 9 Gy + Veh vs 9 Gy + DH) **(*Fig. 12 A1* & *A3*)**. Similar analyses revealed that irradiation increased the levels of glutamate (0 Gy + Veh vs 9 Gy + Veh, p < 0.0001) and DH treatment reversed this effect by reducing the level of glutamate (9 Gy + Veh vs 9 Gy + DH, p = 0.003) ***(Fig. 12 A2)*.**

**Fig. 12:**
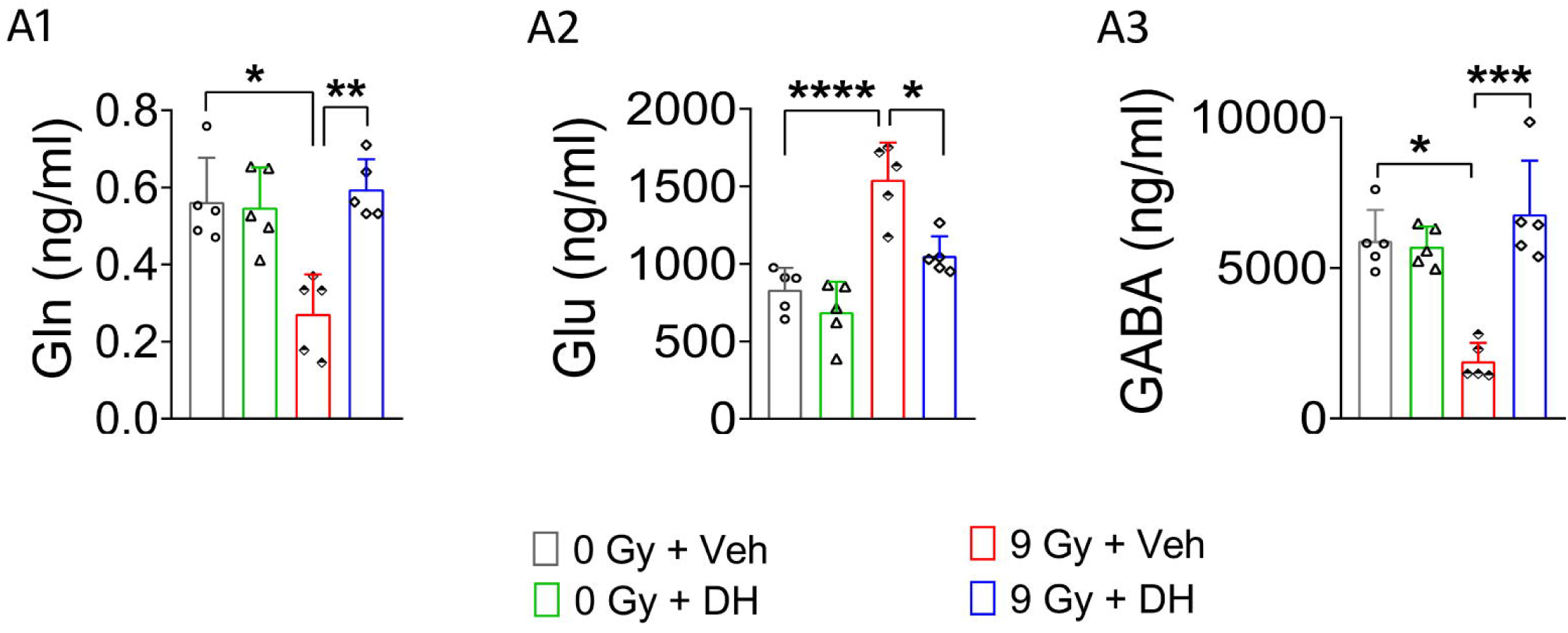
Impaired glutamate-glutamine/GABA cycle in irradiated mice brain: Irradiated mice showed enhanced level of glutamate (A2), and a reduced level of glutamine and GABA (A1 and A3), while DH ameliorates this situation. Two-way ANOVA followed by Tukey’s multiple comparison. ***p < 0.001, ****p < 0.0001. **N** = 5 per group.

## Discussion

This study demonstrates that cranial irradiation induces progressive dysregulation of glucose metabolism and mitochondrial energetics, a disturbed glutamine–glutamate/GABA cycle, and altered glucose transporter expression, potentially contributing to mood and memory impairments and accelerated brain aging. Notably, we provide evidence that oral administration of DH improves mood and memory, regulates glucose metabolism and mitochondrial energetics, and preserves cognitive function in irradiated mice. Furthermore, DH enhances the expression of microglial and neuronal glucose transporters, key regulators of cellular glucose uptake, suggesting a potential mechanism underlying its neuroprotective effects.

Cognitive performance was assessed using NOR, OiP, TO, and WMT paradigms, which revealed selective impairments in hippocampal- and prefrontal cortex–dependent spatial and temporal memory, consistent with prior reports (Kesharwani et al., 2025a; Kesharwani et al., 2025b; Kesharwani, 2026; Kiffer et al., 2019; Parihar et al., 2020). To the best of our knowledge, this study is the first to connect cognitive dysfunction with altered energy metabolism and mitochondrial deficits, highlighting the role of DH as a therapeutic intervention for radiation-induced cognitive deficits via regulation of energy metabolism and mitochondrial energetics. Interestingly, in a diabetic mouse model, DH treatment significantly improved performance in multiple cognitive and depression-related behavioral tasks, including NOR and FST(Kesharwani et al., 2025a; Kesharwani et al., 2025b). Collectively, these findings highlight DH as a promising therapeutic strategy to attenuate cognitive decline across diverse models.

Our findings implicate cranial irradiation in cognitive aging, the precise molecular mechanisms underlying the onset and progression of radiation induced cognitive decline remain incompletely understood. This process is thought to involve changes in the neuroimmune transcriptome, abnormalities in anti-inflammatory and proinflammatory signalling, and difficulties with mitochondrial energy generation (Boyd et al., 2021; Greene-Schloesser et al., 2013; Lumniczky et al., 2017). Our integrated transcriptomic and proteomic analyses revealed widespread dysregulation of genes involved in ATP production and energy metabolism in irradiated mouse brains. Key mitochondrial and metabolic genes (e.g., *mt-Cytb, mt-Nd1, mt-Atp6, mt-Nd2,* and *Kcnj13*) were differentially expressed, indicating impaired oxidative phosphorylation, glucose metabolism, and mitochondrial energetics. Importantly, DH treatment partially restored energy-related pathways by upregulating genes associated with mitochondrial function, neurotransmission, and neuroprotection, while also modulating distinct targets (e.g., *Sgk1, Plekhf1,* and *Pmch*) independent of irradiation effects. Additionally, we found several genes linked to synaptic plasticity, ion channel regulation, and neuroinflammation (e.g., *Oacyl, Kcnj13, Phxr4,* and *Cd6*) were differentially expressed, providing interaction between metabolic dysfunction and immune signaling, suggesting another novel target for radiation induced cognitive deficits.

Further, our study revealed that cranial irradiation potentially alters the key enzymes regulating aerobic glycolysis and energy metabolism (PDH, LDH, α-KGDH, PFKM, G6PDH), were altered, which were positively regulated by the DH treatment in irradiated mice. These findings indicated that disrupted energy metabolism and neuronal energy utilization, and its regulation could be the major target against radiation induced cognitive deficits. These changes likely contribute to energy failure through mechanisms such as blood–brain barrier disruption, reduced glucose transport, and altered metabolic signaling, ultimately impairing synaptic function (Brown and Thore, 2011; Kapogiannis and Mattson, 2011; Monje and Palmer, 2003). Moreover, previous studies have shown that reduced glucose transporter expression leads to energy deprivation, contributing to neurodegeneration, neuroglycopenia, impaired synaptic function, and reduced ATP production (Chen et al., 2023; Han et al., 2021; Marin-Valencia et al., 2012). Inconsistent to this, our study revealed that radiation significantly reduced the neuronal (GLUT3) and microglial (GLUT5) glucose transporter in the hippocampus of the irradiated brain. In addition we noticed, DH increase the cell-specific transporters (GLUT3, and GLUT5), suggesting the maintaining the glucose uptake, managing neurotransmitter levels, and sustaining ion density (Koepsell, 2020). These studies reports that compromised glucose transport, caused by dysfunctional glucose transporters, plays a role in the onset of Alzheimer’s disease (AD) and the repercussions of prolonged stress (Szablewski, 2017). Although our study examined the expression of cell-specific glucose transporters, their functional activity remains to be determined.

Previous studies reported glutamate serves as a vital energy substrate for the brain and is integral to this process, and altered glutamine metabolisms have been associated with mitochondrial dysfunction and energy dysregulation (Nobili et al., 2017). In alignment with this, we observed increased glutamate and glutamine levels alongside reduced GABA in the hippocampus, suggesting that irradiation disrupts the glutamate–glutamine cycle. These metabolic alterations may lead to imbalances in neurotransmission, altered brain activity, and changes in energy demand and glucose utilization. Further, our omics data corroborate the disruption of key genes involved in glutamine metabolism and neurotransmission, including glutamate decarboxylase 2 (GAD2) and glutamate receptor alpha 1 (GLRA1). Genes associated with GABAergic transmission, including GABRQ, ABAT, GABRR2, HTR4, GAD2, and Gabarapl1, exhibit distinct expression patterns within the glutamine–glutamate/GABA cycle in irradiated mice. In the irradiated brain, these coordinated alterations collectively suggest a dysregulation of the excitatory–inhibitory balance, potentially contributing to impaired synaptic signalling, altered neurotransmitter recycling, and compromised neuronal network stability. Moreover, DH treatment, positively regulated the glutamate-glutamine/GABA cycle by regulating at transcription levels of genes critical for glutamine-glutamate homeostasis (*GLRA1*) and GABAergic function (*GAD2*) in irradiated mice.

In summary, our study demonstrates that cranial irradiation-induced disruption of energy metabolism is a key contributor to cognitive decline, and highlighting the metabolic regulation as a potential therapeutic strategy to improve radiation induced cognitive deficits.

## Authors’ contributions

**AK**: Data curation, Formal analysis, Methodology, Software, Validation, Visualization, Writing – original draft, Writing – review & editing. **PB:** Data curation, Methodology, Validation. **AA:** Data curation, Methodology, Validation. **RC:** Data curation, Methodology, investigation, resources **VT:** Data curation, investigation, resources **KP:** Investigation, Software, Writing – review & editing. **VR**: Funding acquisition, Investigation, Project administration, Resources, Software, Supervision. **VKP:** Conceptualization, Funding acquisition, Investigation, Project administration, Resources, Supervision, Validation, Writing – review & editing.

## Availability of data and material

Data generated in this study are included in the manuscript and supplementary information.

## Acknowledgments

We sincerely thank *Dr. Peraman Ramalingam* and *Dr. Rahul Laxman Gajbhiye* from the Department of Pharmaceutical Analysis for providing access to the Liquid Chromatography-High Resolution Mass Spectrometry (LC-HRMS) platform used in the proteomics analysis for this study. Additionally, we extend our gratitude to *ICMR-RMRIMS, Patna,* for providing access to the confocal microscope facility, which was essential for the imaging and analysis performed in this study.

## Abbreviations

DH: Dehydrozingerone
CNS: Central Nervous System
FST: Forced Swim Test
NOR: Novel Object Recognition
OiP: Object in Placement
TO: Temporal Order
WMT: Water Maze Test
mPFC: medial Prefrontal Cortex
PDH: Pyruvate dehydrogenase
LDH: Lactate dehydrogenase
α-KGDH: α-ketoglutarate dehydrogenase
PFKM: Phosphofructokinase
HK: Hexokinase
G6PDH: Glucose 6 Phosphate dehydrogenase
AD: Alzheimer disease
GLUT 3: Glucose transporters 3
GLUT 5: Glucose transporters 5

## References

Acharya, M.M., Green, K.N., Allen, B.D., Najafi, A.R., Syage, A., Minasyan, H., Le, M.T., Kawashita, T., Giedzinski, E., Parihar, V.K., West, B.L., Baulch, J.E., Limoli, C.L., 2016. Elimination of microglia improves cognitive function following cranial irradiation. Sci Rep 6, 31545.

Barker, G.R., Warburton, E.C., 2015. Object-in-place associative recognition memory depends on glutamate receptor neurotransmission within two defined hippocampal-cortical circuits: a critical role for AMPA and NMDA receptors in the hippocampus, perirhinal, and prefrontal cortices. Cereb Cortex 25, 472–481.

Boyd, A., Byrne, S., Middleton, R.J., Banati, R.B., Liu, G.J., 2021. Control of Neuroinflammation through Radiation-Induced Microglial Changes. Cells 10.

Brown, W.R., Thore, C.R., 2011. Review: cerebral microvascular pathology in ageing and neurodegeneration. Neuropathol Appl Neurobiol 37, 56–74.

Chen, Y., Joo, J., Chu, J.M., Chang, R.C., Wong, G.T., 2023. Downregulation of the glucose transporter GLUT 1 in the cerebral microvasculature contributes to postoperative neurocognitive disorders in aged mice. J Neuroinflammation 20, 237.

Christie, L.A., Acharya, M.M., Parihar, V.K., Nguyen, A., Martirosian, V., Limoli, C.L., 2012. Impaired cognitive function and hippocampal neurogenesis following cancer chemotherapy. Clin Cancer Res 18, 1954–1965.

Culhane, A.C., Thioulouse, J., Perrière, G., Higgins, D.G., 2005. MADE4: an R package for multivariate analysis of gene expression data. Bioinformatics 21, 2789–2790.

Dobin, A., Davis, C.A., Schlesinger, F., Drenkow, J., Zaleski, C., Jha, S., Batut, P., Chaisson, M., Gingeras, T.R., 2013. STAR: ultrafast universal RNA-seq aligner. Bioinformatics 29, 15–21.

Greene-Schloesser, D., Moore, E., Robbins, M.E., 2013. Molecular pathways: radiation-induced cognitive impairment. Clin Cancer Res 19, 2294–2300.

Han, R., Liang, J., Zhou, B., 2021. Glucose Metabolic Dysfunction in Neurodegenerative Diseases-New Mechanistic Insights and the Potential of Hypoxia as a Prospective Therapy Targeting Metabolic Reprogramming. Int J Mol Sci 22.

He, B., Bi, K., Jia, Y., Wang, J., Lv, C., Liu, R., Zhao, L., Xu, H., Chen, X., Li, Q., 2013. Rapid analysis of neurotransmitters in rat brain using ultra-fast liquid chromatography and tandem mass spectrometry: application to a comparative study in normal and insomnic rats. J Mass Spectrom 48, 969–978.

Kapogiannis, D., Mattson, M.P., 2011. Disrupted energy metabolism and neuronal circuit dysfunction in cognitive impairment and Alzheimer’s disease. Lancet Neurol 10, 187–198.

Kesharwani, A., Lahamge, D., Singh, S.K., Ravichandiran, V., Parihar, V.K., 2025a. Potentiation of endocannabinoid signaling alleviates depressive-like behavior in diabetic mice. Pharmacol. Res. - Reports 3.

Kesharwani, A., Sree, B.K., Singh, N., Gajbhiye, R.L., Murti, K., Peraman, R., Pandey, K., Limoli, C.L., Velayutham, R., Parihar, V.K., 2025b. Dehydrozingerone Improves Mood and Memory in Diabetic Mice via Modulating Core Neuroimmune Genes and Their Associated Proteins. ACS Pharmacol Transl Sci 8, 1694–1710.

Kesharwani, A.L., D; Sharma, A; Kumarasamy, M; Ravichandiran, V; Parihar, V. K., 2026. Cannabidiol rescues age-associated cognitive decline in mouse model. Bioorxiv.

Kiffer, F., Boerma, M., Allen, A., 2019. Behavioral effects of space radiation: A comprehensive review of animal studies. Life Sciences in Space Research 21, 1–21.

Kodali, M., Parihar, V.K., Hattiangady, B., Mishra, V., Shuai, B., Shetty, A.K., 2015. Resveratrol prevents age-related memory and mood dysfunction with increased hippocampal neurogenesis and microvasculature, and reduced glial activation. Sci Rep 5, 8075.

Koepsell, H., 2020. Glucose transporters in brain in health and disease. Pflugers Archiv : European journal of physiology 472, 1299–1343.

Liao, Y., Smyth, G.K., Shi, W., 2019. The R package Rsubread is easier, faster, cheaper and better for alignment and quantification of RNA sequencing reads. Nucleic Acids Res 47, e47.

Lim, C.H., Soga, T., Parhar, I.S., 2023. Social stress-induced serotonin dysfunction activates spexin in male Nile tilapia (Oreochromis Niloticus). Proc Natl Acad Sci U S A 120, e2117547120.

Lumniczky, K., Szatmári, T., Sáfrány, G., 2017. Ionizing Radiation-Induced Immune and Inflammatory Reactions in the Brain. Frontiers in immunology 8, 517.

Makale, M.T., McDonald, C.R., Hattangadi-Gluth, J.A., Kesari, S., 2017. Mechanisms of radiotherapy-associated cognitive disability in patients with brain tumours. Nat Rev Neurol 13, 52–64.

Marin-Valencia, I., Good, L.B., Ma, Q., Duarte, J., Bottiglieri, T., Sinton, C.M., Heilig, C.W., Pascual, J.M., 2012. Glut1 deficiency (G1D): epilepsy and metabolic dysfunction in a mouse model of the most common human phenotype. Neurobiol Dis 48, 92–101.

Mergenthaler, P., Lindauer, U., Dienel, G.A., Meisel, A., 2013. Sugar for the brain: the role of glucose in physiological and pathological brain function. Trends in neurosciences 36, 587–597.

Mi, H., Huang, X., Muruganujan, A., Tang, H., Mills, C., Kang, D., Thomas, P.D., 2017. PANTHER version 11: expanded annotation data from Gene Ontology and Reactome pathways, and data analysis tool enhancements. Nucleic Acids Res 45, D183–d189.

Monje, M.L., Palmer, T., 2003. Radiation injury and neurogenesis. Curr Opin Neurol 16, 129–134.

Nobili, A., Latagliata, E.C., Viscomi, M.T., Cavallucci, V., Cutuli, D., Giacovazzo, G., Krashia, P., Rizzo, F.R., Marino, R., Federici, M., De Bartolo, P., Aversa, D., Dell’Acqua, M.C., Cordella, A., Sancandi, M., Keller, F., Petrosini, L., Puglisi-Allegra, S., Mercuri, N.B., Coccurello, R., Berretta, N., D’Amelio, M., 2017. Dopamine neuronal loss contributes to memory and reward dysfunction in a model of Alzheimer’s disease. Nat Commun 8, 14727.

Owonikoko, T.K., Arbiser, J., Zelnak, A., Shu, H.K., Shim, H., Robin, A.M., Kalkanis, S.N., Whitsett, T.G., Salhia, B., Tran, N.L., Ryken, T., Moore, M.K., Egan, K.M., Olson, J.J., 2014. Current approaches to the treatment of metastatic brain tumours. Nat Rev Clin Oncol 11, 203–222.

Parihar, V.K., Allen, B., Tran, K.K., Macaraeg, T.G., Chu, E.M., Kwok, S.F., Chmielewski, N.N., Craver, B.M., Baulch, J.E., Acharya, M.M., Cucinotta, F.A., Limoli, C.L., 2015. What happens to your brain on the way to Mars. Sci Adv 1.

Parihar, V.K., Angulo, M.C., Allen, B.D., Syage, A., Usmani, M.T., Passerat de la Chapelle, E., Amin, A.N., Flores, L., Lin, X., Giedzinski, E., Limoli, C.L., 2020. Sex-Specific Cognitive Deficits Following Space Radiation Exposure. Front Behav Neurosci 14, 535885.

Parihar, V.K., Limoli, C.L., 2013. Cranial irradiation compromises neuronal architecture in the hippocampus. Proc Natl Acad Sci U S A 110, 12822–12827.

Parihar, V.K., Syage, A., Flores, L., Lilagan, A., Allen, B.D., Angulo, M.C., Song, J., Smith, S.M., Arechavala, R.J., Giedzinski, E., Limoli, C.L., 2021. The Cannabinoid Receptor 1 Reverse Agonist AM251 Ameliorates Radiation-Induced Cognitive Decrements. Front Cell Neurosci 15, 668286.

Prasad, S.R., Kumar, P., Mandal, S., Mohan, A., Chaurasia, R., Shrivastava, A., Nikhil, P., Aishwarya, D., Ramalingam, P., Gajbhiye, R., Singh, S., Dasgupta, A., Chourasia, M., Ravichandiran, V., Das, P., Mandal, D., 2022. Mechanistic insight into the role of mevalonate kinase by a natural fatty acid-mediated killing of Leishmania donovani. Sci Rep 12, 16453.

Singh, N., Kesharwani, A., Sankar, S.H.H., Gajbhiye, R.L., Peraman, R., Bharathavikru, R.S., Pandey, K., Velayutham, R., Parihar, V.K., 2025. Dehydrozingerone ameliorates renal structure compromised in diabetic nephropathy. Biochim Biophys Acta Mol Basis Dis 1871, 167894.

Szablewski, L., 2017. Glucose Transporters in Brain: In Health and in Alzheimer’s Disease. Journal of Alzheimer’s disease : JAD 55, 1307–1320.

van Oostrum, M., Blok, T.M., Giandomenico, S.L., Tom Dieck, S., Tushev, G., Fürst, N., Langer, J.D., Schuman, E.M., 2023. The proteomic landscape of synaptic diversity across brain regions and cell types. Cell 186, 5411–5427.e5423.

