## Supplementary file 1 for "Dehydrozingerone mitigates energy deficits and cognitive impairments induced by cranial irradiation"

Anuradha Kesharwani^,#^, PavanKalyan Banavath^b*^, Akuthota Akanksha^a*^, Richa Chauhan^c^, Vinita Trivedi^c^, Krishna Pandey^d^, Velayutham Ravichandiran^a,e^, Vipan Kumar Parihar^a,b,@^
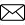

^a^Department of Pharmacology and Toxicology, National Institute of Pharmaceutical Education and Research, Hajipur - 844102, India

^b^Department of Regulatory Toxicology, National Institute of Pharmaceutical Education and Research, Hajipur - 844102, India

^c^Department of Radiation Oncology, Mahavir Cancer Sansthan & Research Centre, Patna- 801505, India

^d^ Department of Clinical Medicine, ICMR-Rajendra Memorial Research Institute of Medical Sciences, Patna 800007, India

^e^Delhi Pharmaceutical Sciences and Research University, New Delhi - 110017, India

* Equal contribution.

^#^ Current affiliation: Florida Atlantic University, Stiles-Nicholson Brain Institute, Jupiter, Florida- 33458, USA

^@^ Current affiliation: Central University of Haryana, Mahendergarh, Haryana-123031, India

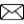
Corresponding Author

Dr. Vipan K. Parihar

Department of Pharmacology and Toxicology,

Department of Regulatory Toxicology,

National Institute of Pharmaceutical Education and Research

Hajipur - 844102, Bihar, India

**Methods**

**Behavioral assessment:**

**Novel Object Recognition (NOR) test:** The Novel Object Recognition (NOR) test was conducted after three days of habituation, during which animals were placed in an open arena for ten minutes daily. Objects with identical material properties but differing in shape and color were used across all behavioral tests. To prevent displacement by the mice, the objects were magnetized and positioned 7 cm from opposing corners and 16 cm apart. During the testing session, mice were exposed to two identical objects for five minutes to familiarize them. Afterward, the animals were returned to their home cages for a 5-minute interval, during which the arena and objects were cleaned with 70% ethanol, and one of the identical objects was replaced with a novel one. Mice were then given an additional five minutes to explore both the familiar and the novel objects. The objects were counterbalanced across groups and excluded from subsequent tests. Trials were scored by an observer blinded to the experimental groups, and results were analyzed using a discrimination index. An exploration score was recorded when the animal approached within 1 cm of an object and directed its gaze towards it. The discrimination index was calculated to evaluate the preference or indifference to novelty (Acharya et al., 2015; Kesharwani et al., 2025a; Kesharwani et al., 2025b; Kesharwani, 2026; Parihar et al., 2015a; Parihar et al., 2016).

**Object in Place (OiP) task:** Following the NOR assessment, mice underwent a two-day habituation period in an open arena, with sessions lasting 10 minutes each day. On the third day, the mice were introduced to four objects of varying shapes, colours, and sizes for 5 minutes to familiarize themselves. After this familiarization period, the mice were returned to their home cages for 5 minutes within the same room. During this time, the arena and objects were cleaned with 70% ethanol, and the positions of two out of the four objects were swapped. All objects were treated uniformly across groups to ensure no bias towards any specific object. The mice were then reintroduced to the arena with the altered object positions for 5 minutes. An observer, blinded to the experimental groups, scored the data, and the results were analyzed using a discrimination index (Barker et al., 2007; Kesharwani et al., 2025a; Kesharwani et al., 2025b; Kesharwani, 2026; Parihar et al., 2015a; Parihar et al., 2016; Parihar et al., 2018; Parihar et al., 2021).

**Temporal Order (TO) task:** After a two-day habituation period in an open arena (10 minutes per day), a temporal test was conducted, consisting of three phases with specific modifications in each. In Phase 1, mice were exposed to two identical objects for 5 minutes. Subsequently, the mice were returned to their home cages for a 4-hour rest period, during which the arena and objects were cleaned with 70% ethanol. Following this rest, the mice were introduced to two new objects of the same shape as in Phase 1 for another 5 minutes. After the second exposure, the mice were returned to their cages for a 1-hour resting period, during which the cleaning process was repeated. In the final phase, the mice were placed in the arena containing one object from each of the previous phases for 5 minutes. Data collection was performed by an observer blinded to the experimental groups, and results were analyzed using the discrimination index. Scores were evaluated according to the established criteria for temporal order (Acharya et al., 2015; Kesharwani et al., 2025a; Kesharwani et al., 2025b; Kesharwani, 2026; Parihar et al., 2015a; Parihar et al., 2016; Parihar et al., 2018; Parihar et al., 2021).

**Forced Swim Test (FST):** The Forced Swim Test (FST) is a widely used behavioral assay to evaluate helplessness or depressive-like behavior in rodents. The test involves placing the animal in a water-filled cylinder and measuring its mobility over a 5 to 8-minute period. Behavior characterized by immobility with the head above water, referred to as "floating behavior," serves as an indicator of depressive-like tendencies. In this experiment, each mouse was placed in a glass beaker with an inner diameter of 10 cm and a depth of 14.5 cm, filled with tap water at a temperature of 18–20°C to a depth of 9.5 cm. Before the test, the water depth was adjusted to ensure the mice could not touch the bottom with their hind paws or tails. The test was conducted in a single 5-minute session, during which immobility and struggling were recorded every minute. Struggling was defined as active movements aimed at escaping the container, while immobility or floating was identified as minimal movements necessary to keep the head above water. At the end of the session, each mouse was gently removed from the water, dried with a towel or dryer, and returned to its home cage. The total immobility duration was calculated for each mouse and used as a measure of depressive-like behavior (Kesharwani et al., 2025a; Kesharwani et al., 2025b; Kesharwani, 2026; Parihar et al., 2011; Parihar and Limoli, 2013; Parihar et al., 2021).

**Water Maze Test (WMT):** Hippocampus-dependent spatial learning and memory were evaluated in irradiated mice using the Water Maze test (WMT), following previously established methods (Hattiangady et al., 2011; Hattiangady and Shetty, 2012)The apparatus consisted of a circular pool (170 cm in diameter, 75 cm in height) filled with room-temperature water to a depth of 35 cm. The water was rendered opaque using non-toxic white paint to prevent visual detection of the submerged platform. Spatial orientation was facilitated by prominent extra-maze cues placed on the surrounding walls.

The pool was divided into four quadrants, and a circular escape platform (15 cm diameter) was positioned in the center of one quadrant, submerged 1 cm below the water surface. Illumination was provided indirectly using a quartz halogen lamp directed toward the ceiling. Mouse behavior was recorded using an overhead camera and analyzed with ANY-maze tracking software.

Training was conducted for seven consecutive days, with one session per day consisting of four acquisition trials. In each trial, mice were released into the pool from pseudo-randomized start positions, facing the wall. Animals were given a maximum of 90 seconds to locate the hidden platform, with a 120-second inter-trial interval. The platform location remained constant throughout the acquisition phase. Upon reaching the platform, mice were allowed to remain there for 30 seconds. If a mouse failed to locate the platform within the allotted time, it was gently guided to it and allowed the same rest period. Measured parameters included escape latency, path length, swim path efficiency (ratio of optimal to actual path), and swim speed, all quantified using ANY-maze software. Learning performance was primarily assessed using mean escape latency, as swim speeds did not differ significantly between groups.

For memory retention, a probe trial was conducted 24 hours after the final training session. During this 45-second test, the platform was removed, and mice were released from the quadrant opposite the previous platform location. Memory performance was assessed by time spent in the target quadrant, latency to first entry into the former platform zone, and number of crossings over the previous platform location (Kodali et al., 2015).

**Tissue Processing for RNA Sequencing, Proteomics, ELISA, Immunohistochemistry, and Neurotransmitter Analysis:** Biological and immunohistochemical evaluations were performed 45 days after cranial irradiation, including 14 days following DH treatment. The subsequent section provides a detailed description of the methods and procedures employed.

**RNA Sequencing:** RNA was extracted from 100 mg of freshly separated mPFC using the RNeasy mini kit (catalog number: 74106). The purity of RNA was checked using a NanoDrop ND-100 Spectrophotometer (NanoDrop Technologies, Wilmington, DE) and a Qubit 4Fluorometer (Thermo Fisher Scientific- IN), and the quality of RNA was confirmed using a Tapestation4200TM (Agilent Technologies, Santa Clara, US). The RNA integrity number (RIN) was calculated by comparing the 28S:18S rRNA ratio, which was found to be 2:1 and had a RIN score greater than 9. Extracted RNA samples were collected, with an initial RNA input of 500 ng, for library construction. The "KAPA mRNA Capture kit" was used to capture mRNA with oligo mag beads, which was subsequently broken down with heat and magnesium. The KAPA RNA Hyper prep kit for Illumina sequencing contains all of the enzymes and buffers required to quickly generate stranded mRNA-Seq libraries.

In a first strand synthesis step, reverse transcriptase and random hexamers are employed to produce cDNA. The cDNA is subsequently converted into double-stranded cDNA by replacing thymine with uracil and adding dAMP to the 3′ ends. To amplify the library, insert fragments with the correct adaptor sequences at both ends and use high-fidelity, low-bias PCR in the adapter ligation procedure, which involves ligating dsDNA adapters with 3′ dTMP overhangs to library fragments. The strand bearing the dUTP marker is not amplified, allowing for strand-specific sequencing. Prepared libraries were sequenced on the IlluminaNextseq 2K to generate 40 million, 2x150bp reads per sample. Up to 75% of the sequenced nucleotides had a Q30 value of more than 90. After the sequencing data was processed and translated into FASTQ files, the data was analyzed. Fastqc was used to detect base quality and contamination from sequencing artifacts, and paired-end raw sequence reads quality reports were generated as a result. Trim Galore was used to trim adapters and sequences of poor quality. The trimmed sequence reads are mapped to the reference transcriptome using the STAR splice aware alignment method. The Rsubread package was used to calculate feature-specific expression counts. The NOISeq R package identifies and eliminates low-count features across samples. To normalize the expression counts, the NOISeq R package's TMM approach was utilized. Significant differential feature estimation was accomplished using the NOISeq R program, which simulates technical duplicates for data without biological replicates. To enrich the functional annotation, the R programme gprofiler2 was employed. A treemap plot and a summary of the Gene Ontology were created using the R package rrvgo (Culhane et al., 2005; Dobin et al., 2013; Kesharwani et al., 2025b; Liao et al., 2019; Singh et al., 2025; Tarazona et al., 2015).

**Tissue proteome analysis using high-resolution mass spectrometry and nano-liquid chromatography:** Proteomic analysis of brain tissue was conducted using a NanoLC (Q-Orbitrap) mass spectrometry instrument. Tissue protein samples were digested with trypsin in 50 mM acetic acid at a 1:50 enzyme-to-protein ratio. The digested samples were vacuum-dried at 30 °C for 1 hour and 30 minutes in AH-L mode using a vacuum concentrator (Concentrator Plus, Eppendorf) and stored at -80 °C until further analysis. Before LC-MS analysis, the dried peptides were reconstituted in 0.1% formic acid. Mass spectrometric analysis was performed using an Orbitrap Exploris 240 mass spectrometer (LTQ-XL, Thermo Fisher Scientific). Peptide fractionation and chromatographic separation were carried out with a Nano-LC system (Easy-nLC 1000) equipped with a C18 Easy-Spray nano column (75 μm ID × 2 cm, 3 μm particle size) at a flow rate of 5 μL/min in 0.1% formic acid. After loading onto the pre-column, peptides were transferred to the analytical C18 column (75 μm ID × 15 cm, 3 μm particle size), which was equilibrated with 95% solvent A (0.1% formic acid) and 5% solvent B (80% acetonitrile, 0.1% formic acid). Peptide elution was performed at a flow rate of 300 nL/min, with a 140-minute gradient of solvent B as follows: 5-15% for 40 min, 15-50% for 45 min, 50-95% for 20 min, 95% for 10 minutes, and 5% for 25 min with a 2 µl injection volume. The mass spectrometer operated in data-dependent acquisition mode with XCalibur software. Survey scans were performed in the Orbitrap within the 350–2000 m/z range at a resolution of 60,000. Ion fragmentation was achieved using collision-induced dissociation (CID). Protein identification and quantification were performed using the full dataset analyzed with Proteome Discoverer (PD; version 2.5.0; Thermo Fisher Scientific)(Kesharwani et al., 2025b; Prasad et al., 2022; Singh et al., 2025).

**ELISA Assay:** The levels of 6-phosphofructokinase (PFKM), glucose-6-phosphate dehydrogenase (G6PDH), hexokinase (HK), pyruvate dehydrogenase (PDH), lactate dehydrogenase (LDH)*,* α-ketoglutarate dehydrogenase (α-KGDH), Pyruvate and Lactate in tissue homogenates of mPFC were quantified using a following the manufacturer’s instructions.

- **Phosphofructokinase (PFKM), Lactate dehydrogenase (LDH) assay:** To measure PFKM levels in the mPFC, an ELISA kit from MyBiosource (Catalogue No. MBS099507) was used following the manufacturer's instructions. All reagents and samples were brought to room temperature (18°C–25°C) naturally before initiating the assay. Hot water baths were not used to avoid compromising the integrity of the reagents or samples. The ELISA plate was removed from the foil pouch, and any unused strips were returned to the pouch with the desiccant pack to prevent moisture exposure. Blank, Standard, and Sample wells were designated as per the protocol. Blank wells were left empty, while 50 µL of Standards (S1–S6) and 50 µL of samples were added to the respective wells. Subsequently, 100 µL of HRP-Conjugate Reagent was added to all wells except the Blank wells. The plate was then covered with a closure membrane and incubated at 37°C for 60 minutes. Following incubation, all wells, including the Blank wells, were washed four times with the provided wash buffer. Next, 50 µL of Chromogen Solution A and 50 µL of Chromogen Solution B were added to each well, with Solution B protected from light exposure. The plate was mixed gently and incubated in the dark at 37°C for 15 minutes. The reaction was stopped by adding 50 µL of Stop Solution to each well. The optical density (O.D.) at 450 nm was measured using an ELISA reader within 15 minutes of adding the Stop Solution, with an optimal reading typically achieved after approximately 5 minutes. This procedure ensured accurate quantification of PFKM levels in the mPFC.
- **Glucose-6-phosphate dehydrogenase (G6PDH) assay:** Subsequently, 1–50 µL of the supernatant was added to duplicate wells of a 96-well plate, and the final volume in each well was adjusted to 50 µL using the G6PDH Assay Buffer. A Master Reaction Mix was prepared, ensuring that 50 µL of the mix was available for each reaction well. Fifty microliters of the Master Reaction Mix were added to the standard, positive control, and sample wells. The plate was protected from light to prevent photochemical reactions, and the contents were mixed thoroughly using a horizontal shaker or by pipetting. After 2–3 minutes, an initial measurement (T_initial_) was taken by recording the absorbance at 450 nm, referred to as (A450)_initial_. It was critical to ensure that (A450)_initial_ fell within the linear range of the standard curve. The plate was incubated at 37 °C, and absorbance measurements were taken every 5 minutes at 450 nm, while keeping the plate protected from light throughout the incubation. Measurements continued until the absorbance of the most active sample approached or exceeded the value of the highest standard (12.5 nmole/well), indicating that the reaction was nearing the end of the linear range of the standard curve. The final measurement, (A450)_final_, for calculating enzyme activity, was determined as the penultimate reading before the most active sample exceeded the linear range. The time corresponding to this measurement was recorded as T_final_. This step ensured accurate quantification of enzyme activity within the linear dynamic range of the assay.
- **Hexokinase (HK) assay:** Tissue samples (0.1 g) were homogenized with 1 ml of assay buffer and centrifuged at 8,000 × g at 4°C for 10 minutes. The supernatant was carefully collected into a new centrifuge tube and stored on ice for subsequent detection. All reagents and their working solutions were prepared according to the manufacturer's specified dilution instructions. Standards were prepared using a two-fold serial dilution method to generate a standard curve. Sample, standard, blank, and positive control wells were designated according to the protocol. The sample, blank, and positive control wells were left empty, while 200 µL of standard solution, 10 µL of sample, and 10 µL of positive control were added to the respective wells. Subsequently, 10 µL of enzyme solution and 180 µL of substrate solution were added to the sample and positive control wells. The mixture was gently mixed, and absorbance was measured at 340 nm at both the 10-second and 130-second time points.
- **Pyruvate dehydrogenase (PDH):** Weigh 0.1 g of tissue, add 1 mL of Extraction Buffer and 10 µL of Reagent II. Homogenize the sample on ice and centrifuge at 600 g for 5 minutes at 4°C. Transfer the supernatant to a new centrifuge tube and discard the pellet. Centrifuge the supernatant again at 11,000 g for 10 minutes at 4°C to separate the supernatant and precipitate. To the precipitate, add 200 µL of Reagent I and 2 µL of Reagent II, resuspend thoroughly, and use it for PDH activity detection. Preheat the microplate reader or visible spectrophotometer for at least 30 minutes, adjusting the wavelength to 605 nm and calibrating the spectrophotometer to zero using deionized water. Incubate the Working Solution at 37°C for mammalian species or 25°C for other species for 10 minutes. In a 96-well plate or micro glass cuvette, add 10 µL of the sample and 190 µL of the Working Solution. Mix well and record the absorbance at 605 nm at 20 seconds (A1) and 1 minute 20 seconds (A2) using the microplate reader. Calculate the change in absorbance (ΔA) as ΔA = A1 - A2.
- **α-Ketoglutarate Dehydrogenase (α-KGDH) activity assay Kit:** Accurately weigh the tissue and add Reagent 1 at a ratio of 1:9 (weight in grams to volume in milliliters). Homogenize the sample in an ice water bath, then centrifuge at 10,000 g for 15 minutes. Collect the supernatant for analysis. Meanwhile, determine the protein concentration of the supernatant. If the supernatant is turbid after centrifugation, repeat the centrifugation until it is clear before use. Prepare a serial dilution of a 0.5 mmol/L standard solution with double-distilled water, using the following recommended concentration gradient: 0, 0.1, 0.2, 0.25, 0.3, 0.35, 0.4, and 0.5 mmol/L. For the standard wells, add 20 µL of the standard solution at various concentrations. For the sample wells, add 20 µL of the sample. For the control wells, add 20 µL of the sample. Next, add 200 µL of working solution to the standard and sample wells, and 200 µL of double-distilled water to the control wells. Then, add 20 µL of Reagent 4 to each well. Mix thoroughly with the microplate reader for 3 seconds and incubate at 37°C for 10 minutes in the dark. After incubation, add 20 µL of Reagent 5 to each well. Mix thoroughly with the microplate reader for 3 seconds and measure the optical density (OD) at 450 nm using the microplate reader.
- **Pyruvate assay:** Tissue samples (0.1 g) were homogenized in 1 mL of assay buffer and centrifuged at 8,000 × g for 10 minutes at 4°C. The resulting supernatant was carefully transferred to a new centrifuge tube and kept on ice for subsequent analysis. All reagents and working solutions were prepared following the manufacturer’s specified dilution instructions. Standards were prepared using a two-fold serial dilution to create a standard curve. Sample, standard, and blank wells were designated as per the protocol. To each well, 75 µL of the sample, standard, or distilled water was added. Subsequently, 25 µL of dye reagent was added to each well, mixed thoroughly, and allowed to stand at room temperature for 2 minutes. Finally, 100 µL of reaction buffer was added to each well with gentle mixing, and the absorbance was recorded at 520 nm using a spectrophotometer.
- **Lactate assay:** Tissue samples (0.1 g) were homogenized in 1 mL of assay buffer and centrifuged at 8,000 × g for 10 minutes at 4°C. The resulting supernatant was carefully transferred to a new centrifuge tube and stored on ice for subsequent analysis. All reagents and working solutions were prepared according to the manufacturer’s instructions. Sample, standard, and blank wells were designated as specified in the protocol. To each well, 60 µL of reaction buffer was added, followed by 20 µL of the sample, standard, or distilled water into their respective wells. Subsequently, 10 µL each of enzyme and coenzyme were added with thorough mixing, and the plate was incubated at room temperature for 5 minutes. Next, 90 µL of dye reagent A and 10 µL of dye reagent B were added to each well, mixed properly, and incubated at room temperature for another 5 minutes. The absorbance was then measured at 450 nm using a spectrophotometer.

**Immunohistochemistry and confocal microscopy:** The day after the final behavioral test, animals were intracardially perfused with saline followed by 4% paraformaldehyde (PFA) in phosphate buffer (pH 7.4). Mice were deeply anesthetized using isoflurane in a plexiglass chamber until respiration ceased. The thoracic cavity was surgically opened, a small incision was made in the right atrium, and normal saline (25 mL, flow rate: 14–16 mL/min) followed by 4% PFA (75 mL, flow rate: 14–16 mL/min) was injected through the left ventricle. The brains were carefully extracted, fixed overnight in 4% PFA, rinsed in phosphate buffer, and immersed in sucrose solutions with increasing concentrations (10% to 30% w/v) until they sank, indicating complete dehydration. The brains were then embedded in cryostat medium, sectioned into 30-µm-thick slices from posterior to anterior, and stored serially in a 24-well plate containing PBS with 0.2% sodium azide.

Three serial sections per hippocampus were selected to evaluate the expression of neuronal and microglial glucose transporters by quantifying GLUT3+ and GLUT5+ cells. Tissue sections were initially blocked in 10% Normal Goat Serum (NGS) and 0.1% Triton X-100 for 30 minutes at room temperature. Subsequently, the sections were incubated overnight at 4°C with primary antibodies (GLUT3: rabbit polyclonal, G-Biosciences, ITT05463; GLUT5: rabbit polyclonal, G-Biosciences, ITT13545) diluted in 2% NGS with 0.1% Triton X-100. After three 5-minute washes with PBS, the sections were incubated for 1 hour at room temperature with secondary antibodies (GLUT3: AF Goat anti-rabbit 594, ThermoFisher, A32740) diluted in 2% NGS.

Additionally, sections were processed for the quantification of neuronal and microglial markers (MAP2+ and IBA1+ cells, respectively). Following a 30-minute blocking step with 10% NGS and 0.1% Triton X-100 at room temperature, the sections were incubated overnight at 4°C with primary antibodies (MAP2: rabbit polyclonal, Abcam, ab254264; IBA1: rabbit polyclonal, G-Biosciences, ITT06442) diluted in 2% NGS containing 0.1% Triton X-100. After washing with PBS (three 5-minute rinses), the sections were incubated with secondary antibodies (MAP2 and IBA1: AF Goat anti-rabbit 488, ThermoFisher, A32731) for 1 hour at room temperature. The sections were counterstained with DAPI (ab228549, Abcam) for 7 minutes, mounted using a slow-fade antifade medium, and analyzed using a laser scanning confocal microscope.

Immunostained sections were imaged using a Carl Zeiss LSM 880 confocal microscope to generate Z-stacks (1024 × 1024 pixels) at 60× magnification, with 1-µm increments covering the entire tissue. Three tissue samples per animal (12 tissues per group) were analyzed. Quantification of Z-stacks, performed under consistent parameters, included assessing glucose transporter expression (GLUT3+ and GLUT5+ cells) and markers of neuronal and microglial populations (MAP2+ and IBA1+ cells) in hippocampal fields. Only staining that met predefined visual criteria was considered for analysis (Acharya et al., 2016; Parihar et al., 2015a; Parihar et al., 2016; Parihar et al., 2015b).

**Preparation of brain samples for neurotransmitter by LC-MS/MS analysis**: The medial prefrontal cortex (mPFC) region of the brain (~50 mg) was carefully dissected on ice and homogenized in 400 µL of methanol. The homogenate was thoroughly mixed with the internal standard (isoprenaline) to ensure uniform blending. The sample was then centrifuged at 12,000 rpm for 10 minutes, and the supernatant was removed. The resulting pellet was vacuum-dried for 2 hours. Subsequently, the dried sample was reconstituted with 200 µL of methanol, mixed thoroughly, and centrifuged again at 12,000 rpm for 5 minutes to obtain a homogeneous solution. The resulting liquid was transferred to sampling vials for analysis using the LC-MS system to measure the levels of glutamine, glutamate, GABA, dopamine, and kynurenic acid, utilizing dedicated standards for each neurotransmitter. Calibration curves for each neurotransmitter were generated using a series of standard solutions at concentrations of 100, 50, 25, 12.5, 6.25, and 3.125 ng/mL.

Neurotransmitter separation was performed using a high-performance liquid chromatography system coupled with a mass spectrometer. A Thermo Hypersil GOLD™ C18 column was used for separation, with a mobile phase consisting of 0.1% formic acid in water (A) and acetonitrile (B), employing a gradient with varying proportions of B in A. The injection volume was set to 20 µL, and the flow rate was maintained at 0.3 mL/min. Detection of analytes was achieved through targeted selective ion monitoring (tSIM) using a heated electrospray ionization (HESI) probe.

The HESI parameters were optimized as follows: the vaporizer temperature was set to 320°C, and the sheath gas, auxiliary gas, and sweep gas flow rates were adjusted to 40, 15, and 1 arbitrary unit, respectively. The positive ion spray voltage was set to 3500 V, and the ion transfer tube temperature was maintained at 300°C. High precision was ensured for tSIM scans by configuring the resolution, microscans, scan width (m/z), and RF lens settings to optimize data acquisition. Data processing and analysis were conducted using Xcalibur Qual Browser version 4.4 (Thermo Fisher Scientific) (He et al., 2013; Lim et al., 2018).

**Reference:**

Acharya, M.M., Green, K.N., Allen, B.D., Najafi, A.R., Syage, A., Minasyan, H., Le, M.T., Kawashita, T., Giedzinski, E., Parihar, V.K., West, B.L., Baulch, J.E., Limoli, C.L., 2016. Elimination of microglia improves cognitive function following cranial irradiation. Sci Rep 6, 31545.

Acharya, M.M., Martirosian, V., Chmielewski, N.N., Hanna, N., Tran, K.K., Liao, A.C., Christie, L.A., Parihar, V.K., Limoli, C.L., 2015. Stem cell transplantation reverses chemotherapy-induced cognitive dysfunction. Cancer Res 75, 676–686.

Barker, G.R., Bird, F., Alexander, V., Warburton, E.C., 2007. Recognition memory for objects, place, and temporal order: a disconnection analysis of the role of the medial prefrontal cortex and perirhinal cortex. J Neurosci 27, 2948–2957.

Culhane, A.C., Thioulouse, J., Perrière, G., Higgins, D.G., 2005. MADE4: an R package for multivariate analysis of gene expression data. Bioinformatics 21, 2789–2790.

Dobin, A., Davis, C.A., Schlesinger, F., Drenkow, J., Zaleski, C., Jha, S., Batut, P., Chaisson, M., Gingeras, T.R., 2013. STAR: ultrafast universal RNA-seq aligner. Bioinformatics 29, 15–21.

Hattiangady, B., Kuruba, R., Shetty, A.K., 2011. Acute Seizures in Old Age Leads to a Greater Loss of CA1 Pyramidal Neurons, an Increased Propensity for Developing Chronic TLE and a Severe Cognitive Dysfunction. Aging Dis 2, 1–17.

Hattiangady, B., Shetty, A.K., 2012. Neural stem cell grafting counteracts hippocampal injury-mediated impairments in mood, memory, and neurogenesis. Stem Cells Transl Med 1, 696–708.

He, B., Bi, K., Jia, Y., Wang, J., Lv, C., Liu, R., Zhao, L., Xu, H., Chen, X., Li, Q., 2013. Rapid analysis of neurotransmitters in rat brain using ultra-fast liquid chromatography and tandem mass spectrometry: application to a comparative study in normal and insomnic rats. J Mass Spectrom 48, 969–978.

Kesharwani, A., Lahamge, D., Singh, S.K., Ravichandiran, V., Parihar, V.K., 2025a. Potentiation of endocannabinoid signaling alleviates depressive-like behavior in diabetic mice. Pharmacol. Res. - Reports 3.

Kesharwani, A., Sree, B.K., Singh, N., Gajbhiye, R.L., Murti, K., Peraman, R., Pandey, K., Limoli, C.L., Velayutham, R., Parihar, V.K., 2025b. Dehydrozingerone Improves Mood and Memory in Diabetic Mice via Modulating Core Neuroimmune Genes and Their Associated Proteins. ACS Pharmacol Transl Sci 8, 1694–1710.

Kesharwani, A.L., D; Sharma, A; Kumarasamy, M; Ravichandiran, V; Parihar, V. K., 2026. Cannabidiol rescues age-associated cognitive decline in mouse model. Bioorxiv.

Kodali, M., Parihar, V.K., Hattiangady, B., Mishra, V., Shuai, B., Shetty, A.K., 2015. Resveratrol prevents age-related memory and mood dysfunction with increased hippocampal neurogenesis and microvasculature, and reduced glial activation. Sci Rep 5, 8075.

Liao, Y., Smyth, G.K., Shi, W., 2019. The R package Rsubread is easier, faster, cheaper and better for alignment and quantification of RNA sequencing reads. Nucleic Acids Res 47, e47.

Lim, S.I., Song, K.H., Yoo, C.H., Woo, D.C., Choe, B.Y., 2018. High-fat diet-induced hyperglutamatergic activation of the hippocampus in mice: A proton magnetic resonance spectroscopy study at 9.4T. Neurochem Int 114, 10–17.

Parihar, V.K., Allen, B., Tran, K.K., Macaraeg, T.G., Chu, E.M., Kwok, S.F., Chmielewski, N.N., Craver, B.M., Baulch, J.E., Acharya, M.M., Cucinotta, F.A., Limoli, C.L., 2015a. What happens to your brain on the way to Mars. Sci Adv 1.

Parihar, V.K., Allen, B.D., Caressi, C., Kwok, S., Chu, E., Tran, K.K., Chmielewski, N.N., Giedzinski, E., Acharya, M.M., Britten, R.A., Baulch, J.E., Limoli, C.L., 2016. Cosmic radiation exposure and persistent cognitive dysfunction. Sci Rep 6, 34774.

Parihar, V.K., Hattiangady, B., Kuruba, R., Shuai, B., Shetty, A.K., 2011. Predictable chronic mild stress improves mood, hippocampal neurogenesis and memory. Mol Psychiatry 16, 171–183.

Parihar, V.K., Limoli, C.L., 2013. Cranial irradiation compromises neuronal architecture in the hippocampus. Proc Natl Acad Sci U S A 110, 12822–12827.

Parihar, V.K., Maroso, M., Syage, A., Allen, B.D., Angulo, M.C., Soltesz, I., Limoli, C.L., 2018. Persistent nature of alterations in cognition and neuronal circuit excitability after exposure to simulated cosmic radiation in mice. Exp Neurol 305, 44–55.

Parihar, V.K., Pasha, J., Tran, K.K., Craver, B.M., Acharya, M.M., Limoli, C.L., 2015b. Persistent changes in neuronal structure and synaptic plasticity caused by proton irradiation. Brain Struct Funct 220, 1161–1171.

Parihar, V.K., Syage, A., Flores, L., Lilagan, A., Allen, B.D., Angulo, M.C., Song, J., Smith, S.M., Arechavala, R.J., Giedzinski, E., Limoli, C.L., 2021. The Cannabinoid Receptor 1 Reverse Agonist AM251 Ameliorates Radiation-Induced Cognitive Decrements. Front Cell Neurosci 15, 668286.

Prasad, S.R., Kumar, P., Mandal, S., Mohan, A., Chaurasia, R., Shrivastava, A., Nikhil, P., Aishwarya, D., Ramalingam, P., Gajbhiye, R., Singh, S., Dasgupta, A., Chourasia, M., Ravichandiran, V., Das, P., Mandal, D., 2022. Mechanistic insight into the role of mevalonate kinase by a natural fatty acid-mediated killing of Leishmania donovani. Sci Rep 12, 16453.

Singh, N., Kesharwani, A., Sankar, S.H.H., Gajbhiye, R.L., Peraman, R., Bharathavikru, R.S., Pandey, K., Velayutham, R., Parihar, V.K., 2025. Dehydrozingerone ameliorates renal structure compromised in diabetic nephropathy. Biochim Biophys Acta Mol Basis Dis 1871, 167894.

Tarazona, S., Furió-Tarí, P., Turrà, D., Pietro, A.D., Nueda, M.J., Ferrer, A., Conesa, A., 2015. Data quality aware analysis of differential expression in RNA-seq with NOISeq R/Bioc package. Nucleic Acids Res 43, e140.
